# METTL3 regulates breast cancer-associated alternative splicing switches

**DOI:** 10.1101/2022.04.11.487899

**Authors:** Cyrinne Achour, Paula Groza, Devi Prasad Bhattarai, Ángel-Carlos Román, Francesca Aguilo

## Abstract

Alternative splicing (AS) enables differential inclusion of exons from a given transcript, thereby contributing to the transcriptome and proteome diversity. Aberrant AS patterns play major roles in the development of different pathologies, including breast cancer. *N*^6^-methyladenosine (m^6^A), the most abundant internal modification of eukaryotic mRNA, influences tumor progression and metastasis of breast cancer, and it has been recently linked to AS regulation. Here, we identify a specific AS signature associated with breast tumorigenesis *in vitro*. We characterize for the first time the role of METTL3 in modulating breast cancer-associated AS programs, expanding the role of the m^6^A-methyltransferase in tumorigenesis. Specifically, we find that both m^6^A deposition in splice site boundaries and in splicing and transcription factor transcripts, such as *PHF5A* and *MYC*, direct AS switches of specific breast cancer-associated transcripts. Finally, we show that five of the AS events validated *in vitro* are associated with a poor overall survival rate for patients with breast cancer, suggesting the use of these AS events as a novel potential prognostic biomarker.

## INTRODUCTION

Alternative splicing (AS) of pre-mRNA is a crucial step in gene expression regulation that enables the coding diversity of the genome by selecting which transcript isoforms are expressed in a cell-specific and temporal manner (1,2). AS results in the differential inclusion of exons that are joined by the spliceosome, a large multi-subunit complex comprised of five small nuclear ribonucleoprotein particles (snRNPs) and numerous proteins (3), yielding to multiple mRNA transcripts for the same given gene. AS is regulated by conserved *cis*-acting RNA elements responsible for the recruitment of splicing factors, which act either as enhancers or as silencers. The splicing outcome is determined by the composition of these RNA regulatory sequences, the differential G/C content between introns and exons, RNA secondary structures and exon/intron lengths (4). In addition, AS is also influenced by chromatin conformation, histone modifications, DNA methylation, and the rate of transcription elongation (5).

AS functions in diverse biological processes including cell growth (6), stem cell-renewal and differentiation (7), and cell death (8), to name a few examples. Despite the advantage to expand cellular function, aberrant AS is typically deleterious to cells and leads to human disease (9). Indeed, recent advances in high-throughput technologies, which have enabled large-scale expression profiling of patient samples, have revealed widespread splicing alterations in both solid tumors and hematologic malignancies (10,11). Splice site mutations and/or dysregulated expression of splicing factors (12,13) result in tumor-associated AS switches *i*.*e*. AS events in neoplastic tissues that are not detected in their normal counterparts. Tumor-associated AS switches have been linked to neoplastic transformation, tumor growth and progression, and resistance to therapy, and therefore can potentially be used as cancer biomarkers or as a tool for developing new-targeted cancer treatments (14).

Breast cancer is the most frequently diagnosed cancer and the leading cause of cancer-related mortality in women worldwide, being metastatic breast cancer incurable with the currently available therapies. Breast cancer is a heterogeneous disease classified into four molecular subtypes based on the presence of hormone receptors: luminal A (progesterone and estrogen receptor positive (PR+/ER+) and human epidermal receptor 2 negative (HER2-), luminal B (PR+/ER+/HER2+), HER2 (PR-/ER-/HER2+), and triple-negative (PR-/ER-/HER2-). Treatment strategies differ according to the molecular subtype. Similar to other human tumors, breast cancer exhibits aberrant AS events due to mutations either within the splicing regulatory elements or at the splice sites of tumor suppressor genes, or dysregulated expression of the splicing machinery (15-20). Additionally, several studies have shown that *MYC* hyperactivation, a common feature in many human cancers, leads to transcriptional upregulation of splicing factors that direct breast cancer-associated AS switches promoting a malignant phenotype (21-23).

Similar to DNA and proteins, RNAs are also substrates for chemical modifications (24). *N*^6^-methyladenosine (m^6^A), the most abundant internal modification in eukaryotic mRNA, has been shown to influence AS (25-28). m^6^A is co-transcriptionally deposited by the methyltransferase-like 3 (METTL3) and METTL14 methyltransferase complex, which partially localizes to nuclear speckles, where splicing occurs (29-31). It has been shown that depletion of the *Drosophila* METTL3 methyltransferase homologue, results in altered AS patterns that influence sex determination (32-34). In addition, depletion of METTL3 led to an m^6^A-dependent RNA structural remodeling that alters the accessibility to m^6^A-binding proteins, affecting the recruitment of the splicing factor hnRNPC, and thereby influencing AS (35). Indeed, hnRNPC has been recently reported to regulate AS in pancreatic ductal adenocarcinoma and non–small cell lung cancer (36,37). Another mechanism by which m^6^A regulates splicing is through the m^6^A reader YTHDC1 (25). YTHDC1 binds to m^6^A-modified mRNA and recruits the splicing factor SRSF3, which promotes exon inclusion, but impedes the binding of SRSF10, which facilitates exon skipping. Moreover, increased m^6^A levels upon depletion of the eraser FTO promoted binding of SRSF2 resulting in exon inclusion in mouse preadipocytes (38). However, an opposite trend was observed in a different cellular context. Specifically, in HEK293T cells another study showed that *FTO* knockout resulted in changes in splicing with exon skipping events being the most prevalent (39). Although the function of m^6^A in AS has been questioned (40), it has been recently shown that deposition of m^6^A near splice junctions positively affect RNA splicing kinetics and modulates hnRNPG binding, an m^6^A reader which influences RNA polymerase II (RNAPII) occupancy patterns and promotes exon inclusion (41,42).

The last decade has unraveled multiple associations of m^6^A modification with different aspects of breast tumorigenesis (43). However, it is still unclear whether this chemical mark contributes to tumor suppression or promotes oncogenicity (44). For instance, studies on METTL3 have revealed that it is overexpressed in breast cancer compared to normal mammary tissues, and its silencing in different breast cancer cell lines has been associated with increased apoptosis and decreased proliferation (45,46). On the contrary, another study reported that not only METTL3 but also other members of the writer complex such as METTL14 and WTAP are downregulated in breast cancer, suggesting that lower levels of m^6^A may contribute to breast tumorigenesis (47). Similar contradictory findings are observed for other players of m^6^A modification, being writers, erasers or readers of m^6^A up- or down-regulated depending on the cellular context (43,44). Mechanistically, m^6^A may dictate the fate of tumor suppressor or oncogenic transcripts (e.g., *BCL2, BNIP3, c-MYC, CXCR4*, and *CYP1B1*), influence the treatment outcomes (e.g., resistance to tamoxifen or doxorubicin *via* methylation of AK4 or miRNA-221–3p) or regulate the stability of pluripotency factors (e.g., *Nanog* and *KLF4*), thus facilitating epithelial-mesenchymal transition (EMT), metastatic progression or the breast cancer stem cell phenotype, among others. Despite the plethora of information showing the implications of m^6^A in breast cancer, the biological relevance of m^6^A in breast tumor-associated AS switches is currently unexplored.

In this study, we identify an AS signature associated with the acquisition of the malignant phenotype of breast cancer *in vitro*. We describe that METTL3 regulates breast cancer-associated AS switches through a direct mechanism involving m^6^A deposition at the proximity of splice sites. Additionally, our data suggests indirect mechanisms by which METTL3 modulates AS in breast cancer through m^6^A deposition on splicing factors and transcriptional regulators of splicing factors such as *PHF5A* and *MYC*, respectively. Notably, our analyses reveal that m^6^A deposition correlates with intronic regions and depletion of METTL3 results in more exon inclusion for specific genes. Finally, we show that five of the *in vitro* validated AS events are associated with a worse prognosis in breast cancer patients, suggesting the use of these AS events as potential prognostic biomarkers.

## MATERIALS & METHODS

### Antibodies

The following commercially available antibodies were used at the indicated concentrations for western blot: Anti-METTL3 (Abcam, ab221795, 1:5,000), Anti-Actin (Sigma, A5441, 1:5,000), Goat Anti-Mouse IgG H&L (HRP) (Abcam, ab6789 1:10,000), Goat Anti-Rabbit IgG H&L (HRP) (Abcam, ab6721, 1:10,000).

### Cell culture

HEK293T, MCF7 and MDA-MB-231 cell lines were cultured in Dulbecco’s Modified Eagle Medium (DMEM, Gibco) supplemented with 10% fetal bovine serum (FBS, Gibco), and 1% penicillin/streptomycin (Gibco). For MCF7 and MDA-MB-231, media was additionally supplemented with 10 µg/ml human insulin (Sigma-Aldrich). MCF10-A cell line was cultured in DMEM/F12 (Sigma-Aldrich) supplemented with 5% heat-inactivated horse serum (Gibco), 20 ng/ml epidermal growth factor (Sigma-Aldrich), 0.5 mg/ml hydrocortisone (Sigma-Aldrich), 100 ng/ml cholera toxin (Sigma-Aldrich), 10 µg/ml insulin (Sigma-Aldrich), and 1% penicillin/streptomycin (Gibco). Cells were cultured at 37°C in a humidified incubator at 5% CO _2_.

### Lentiviruses production and generation of *METTL3* knockdown cell lines

To generate lentiviral particles, HEK293T cells were co-transfected with pLKO.1-Puro containing shRNA1 and shRNA2 against *METTL3* or scramble control (**Supplementary Table 1**), the packaging vector pCMV-dR8.2-dvpr and the envelope vector pCMV-VSV-G (ratio 6:8:2), with Jet-PEI Polyplus following the manufacturer’s instructions. Lentiviral particles were collected after 48 and 72 hours, filtered through a 0.45 µm filter and concentrated using Amicon Ultra-15 Centrifugal Filter (Merck). Knockdown of *METTL3* was obtained by lentiviral transduction with the lentiviral particles in media supplemented with Polybrene (8 µg/ml). Transduced cells were selected by supplementing the culture media with puromycin (1 µg/ml) for an additional 4 days. The efficiency of *METTL3* knockdown was further evaluated by RT-qPCR and western blot analysis.

### Reverse transcription followed by PCR (RT-PCR) and quantitative PCR (RT-qPCR)

Total RNA was extracted using the RNeasy Mini Kit (Qiagen) following the manufacturer’s recommendations. 1 μg of total RNA was reverse transcribed into cDNA using the RevertAid First Strand cDNA Synthesis kit (Invitrogen). Afterwards, PCR was performed using DreamTaq master mix (Thermo Fisher Scientific) for RT-PCRs. Quantitative PCR (qPCR) was performed using the Power Up SYBR Green qPCR Master Mix (Applied Biosystems) using an Agilent Biosystems instrument. *GAPDH* and *βactin* were used as loading control for RT-PCRs and RT-qPCRs, respectively. Primers are described in **Supplementary Table 1**.

### mRNA purification

mRNA was purified using Dynabeads™ following the manufacturer’s recommendations. mRNA was eluted twice with RNase-free water.

### mRNA mass spectrometry analysis

Purified mRNA (100 ng) was analyzed by liquid chromatography-tandem mass spectrometry (LC-MS/MS) at the Proteomics and Modomics core facility, Norwegian University of Science and Technology (NTNU), Norway.

### RNA immunoprecipitation of m^6^A modified transcripts (MeRIP)

m^6^A modified transcripts were immunoprecipitated as described previously (48). Briefly, 5 µg of mRNA was fragmented by using RNA fragmentation reagents (Invitrogen) prior to overnight ethanol precipitation. The fragmented mRNA was recovered by centrifugation at 14,000 rpm and the pellets were resuspended in DEPC water and 10% of the volume used as the input. The remaining fragmented mRNA was then diluted with 100 µl of 5X IP buffer (250 mM Tris pH 7.4, 500 mM NaCl, 0.25% NP-40) and incubated with 10 µg of m^6^A antibody (Abcam, ab151230) in the presence of RNase inhibitors, for 3 hours at 4°C. 30 µl of prewashed Surebeads Protein A magnetic beads (Bio-Rad) were added and incubated for 2 hours at 4°C. Beads were then washed twice with high-salt IP buffer (50 mM Tris pH 7.4, 1 M NaCl, 1 mM EDTA, 1% NP-40), twice with 1X IP buffer and finally once with high-salt IP buffer. The immunoprecipitated RNA was eluted in PK buffer (100 mM Tris-HCl pH 7.5, 50 mM NaCl, 10 mM EDTA) in the presence of Proteinase K (Invitrogen) recovered with Phenol:Chloroform. The input RNA and the immunoprecipitated RNA were subjected to reverse transcription using the VILO Superscript (Invitrogen™) according to the manufacturer’s instructions, followed by qPCR. Primers used for RT-qPCRs are described in **Supplementary Table 1**.

### RNA-seq and differential gene expression analysis

RNA-seq library preparation was carried out at Novogene facilities (https://en.novogene.com/) and sequenced using Illumina HiSeq 2500 platform (Illumina) as 150 bp pair-ended reads. FASTQ reads were pseudoaligned to the human hg38 transcriptome and quantified using Salmon (49). Thereafter, differentially expressed genes (DEG) were obtained using a MATLAB function with a test under the assumption of a negative binomial distribution where the variance is linked to the mean *via* a locally-regressed smooth function of the mean (50). Afterwards, *P-values* were adjusted by estimation of the false discovery rate for multiple hypotheses (51). We only considered the transcripts with reads in at least half of the samples analyzed.

### AS analysis using RNA-seq

To quantify the AS differences between sets of samples we employed the SUPPA2 pipeline (52). Specifically, the Salmon output files generated for the RNA-seq were adapted for the SUPPA2. Splicing events in the human genome were obtained using a specific SUPPA2 script from the human GTF genome hg38 file. Thereafter, the percentage of splicing inclusion (PSI) values for each event were obtained for each sample, and the differential PSI values (ΔPSI) for each condition was calculated along with a *P-value* for each event. *Ad hoc* MATLAB functions were designed to quantify and represent the different analyses from the final SUPPA2 output files. In the case of publicly available datasets (MYC: GSE196325; PHF5A: SRP137027), the same pipeline from FASTQ reads was performed.

### m^6^A and PTC data analysis from public datasets and comparison with AS

The different m^6^A datasets used in the studies (MCF7: GSE143441; MDA-MB-231: GSM5616175; HEK293T: GSE114543) were standardized for comparison. Specifically, they were converted into hg38 and BED format and then subjected to MACS2 for peak detection (53). Afterwards, we compared their results against AS (exon skipping) datasets by a set of scripts that require PERL and Bedtools (54). Fisher’s exact test was applied to assess the statistical significance for the presence of intronic m^6^A sites in significantly spliced exons compared to non-significantly spliced genes. In the case of premature termination codons (PTCs), the splicing events in transcripts annotated as nonsense-mediated decay were analyzed by Fisher’s exact test in a similar manner than in the case of m^6^A.

### *De novo* motif search

m^6^A peaks that were located within flanking introns of a differentially skipped exon were selected. Then the sequence (+/-150 nt) of these peaks was submitted to *de novo* motif search using HOMER (55). Afterwards, random genomic regions with similar properties of these peaks were retrieved for direct comparison of density distribution along the m^6^A region.

### Gene Ontology (GO) analysis

Gene ontology (GO) analysis was performed using the web tool The Database for Annotation, Visualization and Integrated Discovery (DAVID) (https://david.ncifcrf.gov/).

### Analysis of TCGA datasets using SpliceSeq database

A set of validated splicing events was selected and multiple data associated with breast cancer datasets were retrieved from TCGA using SpliceSeq (56). In order to combine the data from different splicing events, a similar approach was used, as previously described (57). Briefly, MATLAB functions were designed to calculate new coefficients for each event using the Lasso function. In addition, the clinical data linked to these datasets using the TCGA portal was characterized. Finally, MATLAB was again used to generate box plots as well as Kaplan-Meier survival curves; statistical *P-values* for every event and clinical feature was also calculated.

### Statistical analysis

Data are shown as mean ± SEM GraphPad Prism version 8.0.0 was used to perform the statistical analysis. The significance was determined using Student *t*-test or Bonferroni test. Probability values of * *P-value* < 0.05, ** *P-value* < 0.01, *** *P-value* < 0.001, **** *P-value* < 0.0001 were considered as statistically significant.

## RESULTS

### Identification of AS events in non-tumorigenic and breast cancer cell lines

To identify genome-wide differential AS events (DSE) occurring during the acquisition of the breast cancer phenotype, we performed RNA-sequencing (RNA-seq) on a breast non-tumorigenic cell line (MCF10-A), and the commonly used luminal A (MCF7) and triple negative (MDA-MB-231) breast cancer models. Reads were then mapped to exon-splice junction sites to determine DSE, including skipped exons (SE), retained introns (RI), mutually exclusive exons (MX), alternative first or last exons (AF or AL), and alternative 5’
s or 3’ splice sites (A5 or A3) (**Supplementary Figure 1A**). The differences of AS isoforms between the breast cancer cell lines and the non-tumorigenic MCF10-A cells were assessed by calculating the change in percent splicing inclusion (ΔPSI) (**Supplementary Table 2**) (52). We identified in total 37680, 37038 and 36514 DSE, corresponding to 14009, 14530 and 14311 genes in the MCF10-A, MCF7 and MDA-MB-231 cell lines, respectively (false discovery rate [FDR] < 0.05; **Figure 1A**). Despite the majority of the DSE (27260) being shared across the three cell lines, the comparison revealed AS events that were unique to the breast cancer cell lines MCF7 and MDA-MB-231; AF and SE being the most represented categories (**Figure 1B-C**). In addition, the PSI values for both MCF7 and MDA-MB-231 had a uniform distribution between enhanced and repressed splice junctions (**Supplementary Figure 1B-C**). We further performed Gene Ontology (GO) and Kyoto Encyclopedia of Genes and Genome (KEGG) pathway enrichment analysis for the DSE in MCF7 and MDA-MB-231. Only few genes (< 50) were enriched in the GO biological process or KEGG pathway, including the terms of “mRNA splicing, *via* spliceosome”, “cell-cell adhesion” and “MAPK signaling cascade”, amongst others (**Figure 1D**). Yet, the DSE in breast cancer cell lines were significantly enriched in the “nucleoplasm”, “cytosol”, “cytoplasm” and “nucleus” terms for the cellular components categories and enriched in the “protein binding and poly(A) RNA binding” category for the molecular function (**Supplementary Figure 1D**).

**Figure 1.**
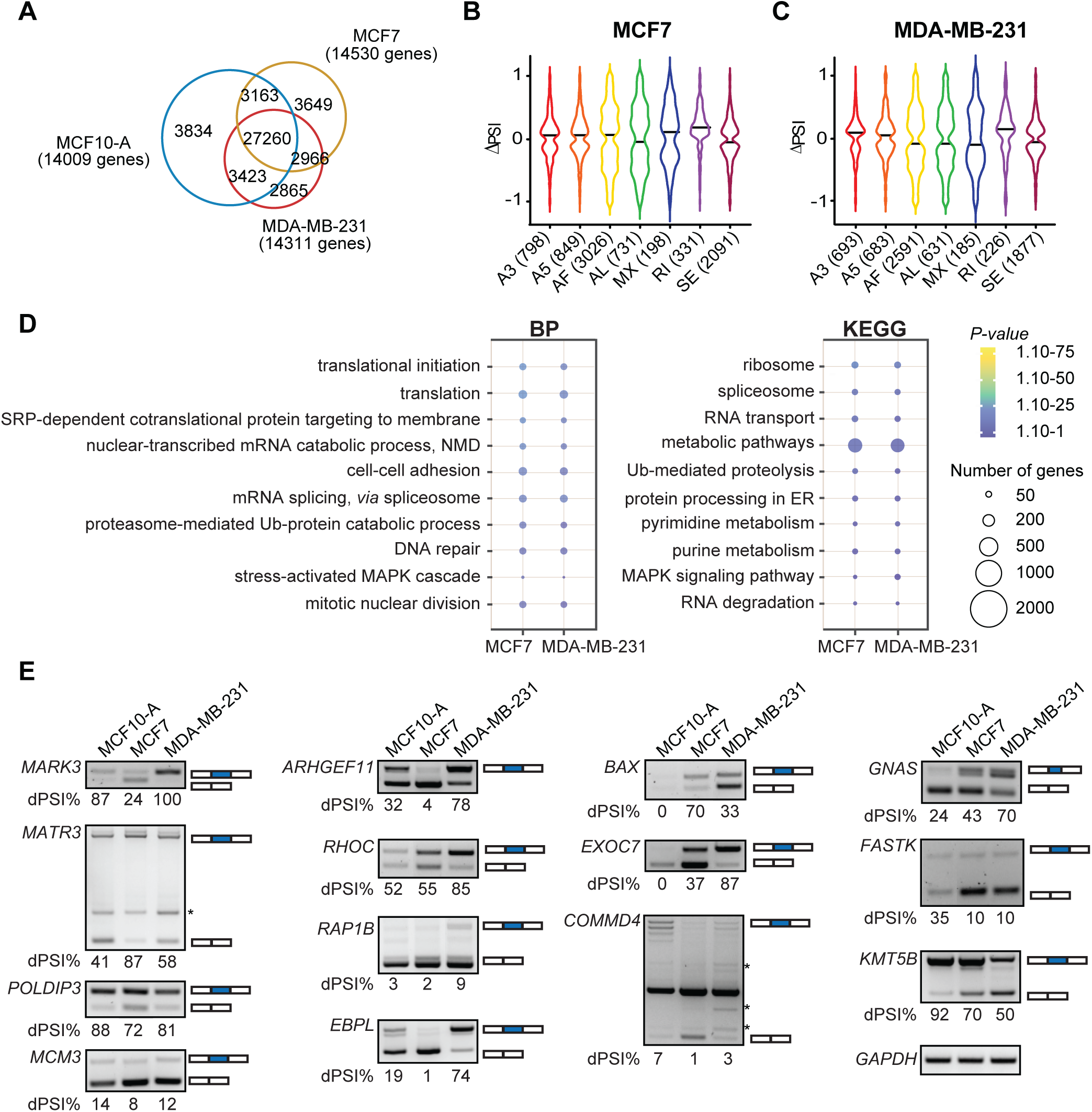
Identification of a breast cancer-associated AS signature. **(A)** Venn diagram showing the number of common AS events and genes in the non-tumorigenic mammary epithelial cell line MCF10-A and the breast cancer cell lines MCF7 and MDA-MB-231. **(B-C)** Violin plots of changes of the significant percent splicing inclusion (△PSI) in the breast cancer cell lines MCF7 and MDA-MB-231 related to the normal mammary epithelial cell line MCF10-A. **(D)** Dot plots representing the GO analysis of the common spliced genes in MCF7 and MDA-MB-231. BP: biological process, KEGG: KEGG pathways. The size and the color of the dots are proportional to the number of genes enriched in each GO term and the significance of the enrichment (1.10^−75^ < *P-value* < 1.10^−1^), respectively. **(E)** RT-PCR showing the DSE between the non-tumorigenic MCF10-A and breast cancer MCF7 and MDA-MB-231 cell lines. The PSI was calculated in percentage for each gene. Non-specific bands are indicated with an asterisk. *GAPDH* was used as a loading control.

We next validated selected AS from genes that were previously associated with different aspects of tumorigenesis (**Figure 1E** and **Supplementary Table 3**). Upon validation of DSE, we observed that *MARK3, MATR3, POLDIP3*, and *MCM3* displayed similar patterns between MCF10-A and MDA-MB-231 cell lines. For *MARK3* and *POLDIP3*, the skipped exon isoform was present at a higher level than the inclusive isoform in MCF7 compared to MCF10-A and MDA-MB-231, whilst the skipped exon isoform for *MATR3* was barely detected in MCF7 cell line. MDA-MB-231 cells displayed distinct AS patterns for *ARHGEF11, RHOC, RAP1B, EBPL, BAX, EXOC7*, and *COMMD4*, whilst AS patterns for *GNAS, FASTK*, and *KMT5B* were similar between MCF7 and MDA-MB-231 but different in comparison to

MCF10-A. There were transcript isoforms showing a positive exon inclusion index, which correlated to a significant negative skipping index of the same exon (e.g. *MARK3, RHOC* and *EBPL*). However, this was not the case for all the validated events (e.g. *EXOC7* and *RAP1B*), suggesting that multiple alternative exons can be spliced in a complex manner (**Figure 1E**). Intriguingly, in the case of *BAX*, we were not able to amplify any isoform for the non-tumorigenic cell line, however we observed that the inclusive isoform was more expressed in MCF7 compared to MDA-MB-231, for which the expression of the skipping exon isoform was higher. Collectively, we have uncovered an AS signature occurring during the acquisition of the malignant phenotype of breast cancer *in vitro*.

### METTL3 regulates AS in breast cancer cell lines

Deposition of m^6^A, catalyzed by METTL3, modulates nearly every aspect of the mRNA lifecycle, including AS (30,58). To determine whether m^6^A regulates AS in breast cancer, we first assessed the expression of METTL3 across non-tumorigenic and breast cancer cell lines. MCF7 and MDA-MB-231 exhibited increased METTL3 expression compared to the non-tumorigenic cell line MCF10-A (**Figure 2A**). Likewise, m^6^A levels on mRNA were also higher in MCF7 and MDA-MB-231 cells (**Figure 2B**). We next performed RNA-seq upon silencing of *METTL3* in MCF10-A, MCF7 and MDA-MB-231 cells. We employed two distinct short-hairpin RNAs (shRNAs) against *METTL3* (sh1 and sh2) to ensure that the observed phenotype is not due to shRNA off-target effects (**Figure 2C-E**). As explained above, reads were mapped to exon-splice junction sites to identify DSE, and both datasets, from sh1 and sh2, were combined to obtain the most significant AS events (**Supplementary Figure 2A-B**; **Supplementary Table 4**). Upon silencing of *METTL3*, we identified 1679 DSE (1072 genes), 2986 DSE (1706 genes) and 3041 DSE (1058 genes) in MCF10-A, MCF7 and MDA-MB-231, respectively (false discovery rate [FDR] < 0.05; **Figure 2F**). Thus, METTL3 depletion accompanied broader modulations in the AS landscape of breast cancer cells lines compared to the non-tumorigenic MCF10-A, suggesting a critical role of METTL3 in regulating tumor-associated AS switches (**Supplementary Figure 2C**). Noteworthy, alterations in DSE upon *METTL3* knockdown were not due to transcriptional changes as gene expression levels were not correlated to ΔPSI (**Supplementary Figure 2D-E**). GO analysis revealed that the common METTL3-regulated AS events in breast cancer cell lines were enriched for the terms “translation”, “regulation of apoptotic process” and “regulation of growth”, suggesting that METTL3 may affect breast tumorigenesis through AS regulation (**Supplementary Figure 2F-G**). Strikingly, GO categories related to “splicing” and “alternative splicing” were highly represented. All types of AS events were affected upon knockdown of *METTL3*, most of the events corresponding to AF in both MCF7 (810 DSE) and MDA-MB-231 (776 DSE) (**Figure 2G-H**). To assess the functional impact of METTL3 in breast cancer, we next performed GO analysis of the differentially expressed genes (DEG) and the DSE of *METTL3* knockdown in MCF7 and MDA-MB-231 cell lines (**Figure 2I-J**). Biological processes frequently altered during tumor progression and metastasis were amongst the most enriched terms. In particular, we observed a common significant enrichment in GO terms associated with “cell adhesion” in both cell lines, and “MAPK cascade” and “apoptosis” in MCF7 and MDA-MB-231, respectively (**Figure 2I-J**). Additionally, we found that the terms “mRNA stability”, “mRNA splicing” and “RNA processing” were specifically enriched in both MCF7 and MDA-MB-231 for the DSE, but these terms were not found among genes whose mRNA levels were affected by *METTL3* silencing. Overall, this data supports the idea that m^6^A may regulate breast tumorigenesis by influencing multiple pathways, including AS of splicing factors and other RNA-binding proteins.

**Figure 2.**
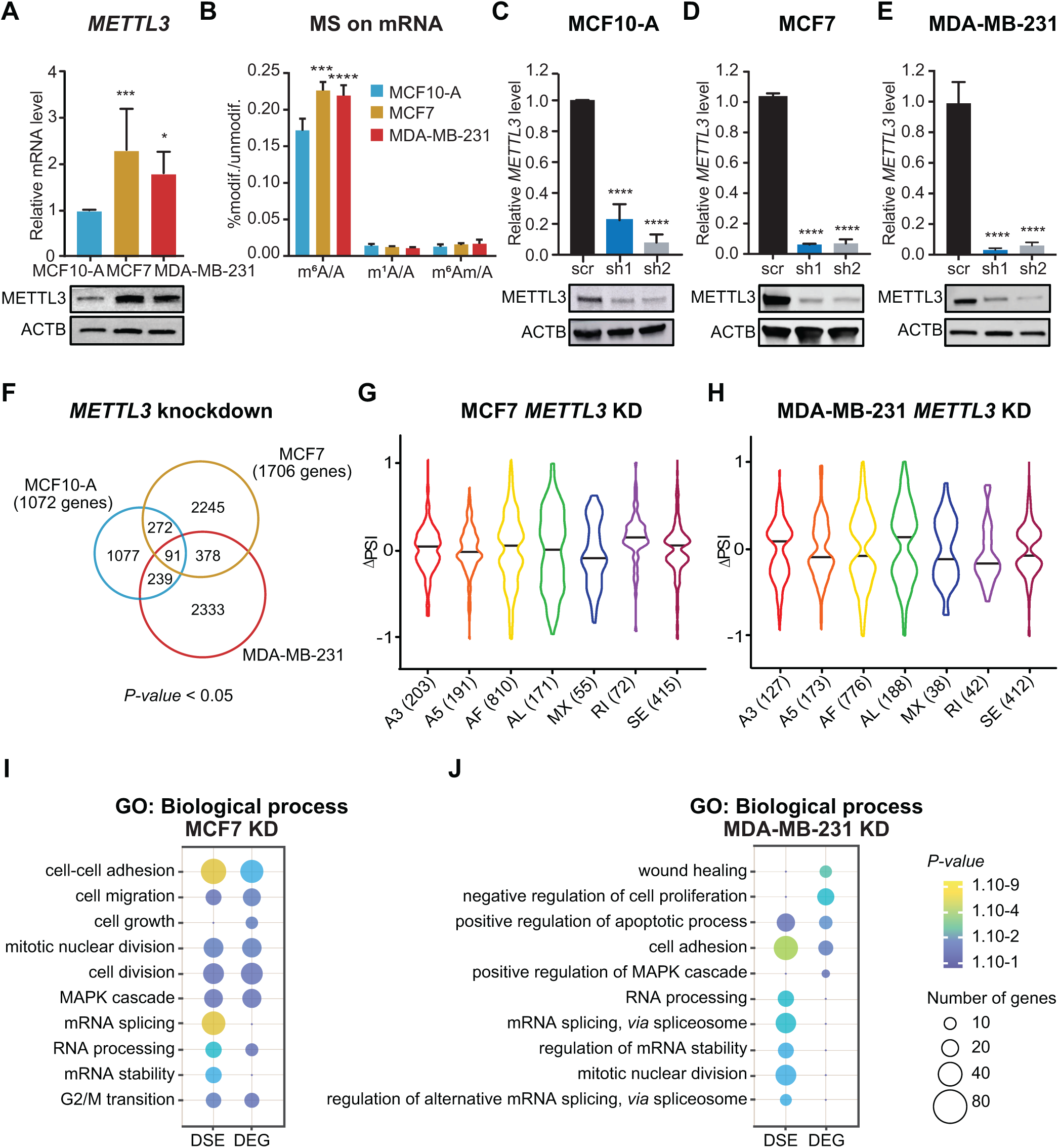
METTL3 regulates AS in breast cancer cell lines. **(A)** RT-qPCR analysis of *METTL3* mRNA level (upper) in the non-tumorigenic mammary epithelial cell line MCF10-A and the breast cancer cell lines MCF7 and MDA-MB-231. *METTL3* is normalized to *β-ACTIN*. n = 3, *** *P-value* < 0.001, * *P-value* < 0.05. Western blot of METTL3 (lower) on whole cell extracts (WCE) from MCF10-A, MCF7 and MDA-MB-231 cell lines. βACTIN (ACTB) is used as the loading control. **(B)** LC-MS/MS quantification of m^6^A, m^1^A and m^6^A _m_ in mRNA of MCF10-A, MCF7 and MDA-MB-231 cell lines. Methylated adenosines are normalized to the total of unmodified adenosines. n = 3; **** *P-value* < 0.0001, *** *P-value* < 0.001. **(C-E)** RT-qPCR analysis of *METTL3* mRNA level in *METTL3* knockdown (sh1 and sh2) and scramble (scr) control in MCF10-A, MCF7 and MDA-MB-231 cell lines. *METTL3* is normalized to *β-ACTIN*. n = 3, *** *P-value* < 0.001, * *P-value* < 0.05. **(F)** Venn diagram showing the common DSE between knockdowns of *METTL3* in MCF10-A (blue), MCF7 (yellow) and MDA-MB-231 (red). The corresponding total number genes is indicated in brackets; *P-value* < 0.05. **(G-H)** Violin plots of the significant percent splicing inclusion (△PSI) in knockdowns of *METTL3* in MCF7 and MDA-MB-231 cells related to control cells. **(I-J)** Dot plots of the GO enrichment analysis of the DSE and the differentially expressed genes (DEG) in MCF7 and MDA-MB-231 upon depletion of METTL3. The size and the color of the dots are proportional to the number of genes enriched in each GO term and to the significance of the enrichment (1.10^−9^ < *P-value* < 1.10^−1^), respectively.

### m^6^A modification regulates AS through several mechanisms

To identify whether METTL3 modulates AS through m^6^A deposition at the proximity of splice sites, we analyzed available m^6^A-RNA immunoprecipitation sequencing (MeRIP-seq) data from chromatin-associated RNAs in HEK293T cells. Although some of the transcripts that underwent DSE upon *METTL3* knockdown harbored the m^6^A mark at exon-intron junction sequences (e.g. *DLG5, LARGE1, INO80C*), we could not detect any significant correlation between intronic m^6^A deposition and AS (**Figure 3A**). Using previously published available data, we next examined the distribution of m^6^A sites at the exon-intron junction in MCF7 and MDA-MB-231 cell lines in total mRNA (59). We found a significant enrichment between m^6^A deposition at exon-intron junction boundaries and processing efficiency (**Figure 3A** and **Supplementary Figure 3A-B**). Notably, more than half of the DSE that we identified in MCF7 and MDA-MB-231 harbor m^6^A modification (**Supplementary Figure 3C**). Sequence logo analysis of both datasets revealed the presence of highly enriched non-DRACH (D=A, G, U; R=A, G; H=A, C, U) motifs in addition to the less enriched m^6^A consensus motif DRACH in the regions ±150 nt around the m^6^A peak summit compared to randomly generated 300 nt genomic intervals (**Figure 3B-C** and **Supplementary Figure 3D**). Altogether, these results suggest that m^6^A directly regulates AS in our cellular models, and that the DSE are cell-type specific. Moreover, these isoforms had coding potential as they were not enriched for PTCs, stop codons that occur >50 nucleotides upstream of the splice junction (60), that would result in nonsense-mediated decay (**Figure 3A**). We next performed GO analysis of the m6A datasets for MCF7 and MDA-MB-231 and observed that again “mRNA and RNA splicing” were amongst the most enriched terms (**Supplementary Figure 3E**). Similar to METTL3-dependent AS switches (**Figure 2I-J**), m^6^A deposition was also prominent in categories important for breast cancer progression and metastasis.

**Figure 3.**
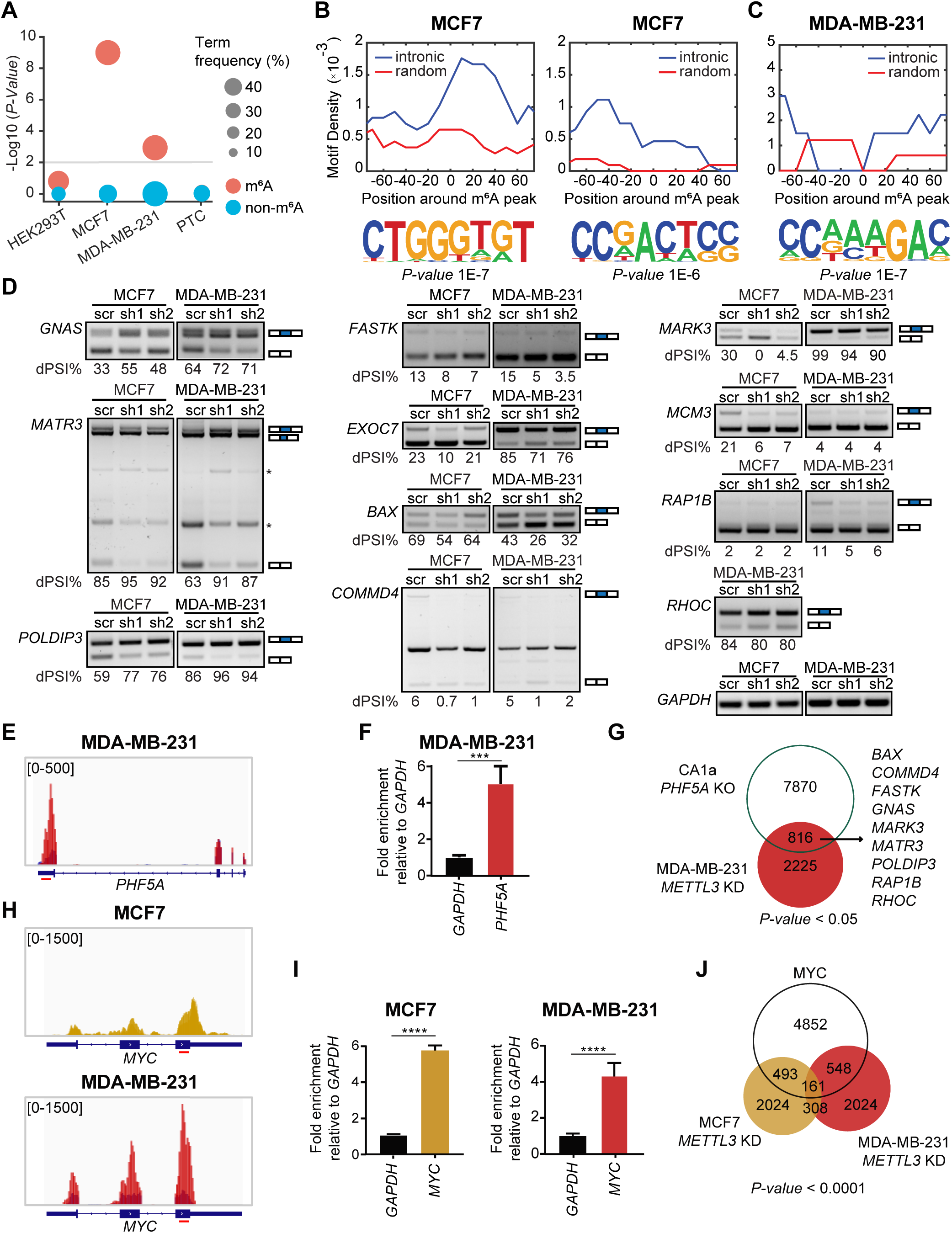
m^6^A influences AS via intronic deposition and through regulation of splicing factors. **(A)** Dot plot representing the level of statistical significance (-Log10 (*P-value*)) of AS transcripts harbouring m^6^A (red) or non-m^6^A modified (blue) in HEK293T (chromatin-bound transcripts), MCF7, MDA-MB-231, and transcripts with premature termination codons (PTC) (60). The size of the dots is proportional to the frequency of the events; the grey line denotes *P-value* < 0.01. **(B-C)** Motif density in the –80 to +80 nt region around the m^6^A peak in intronic (blue) or random regions (red) (upper panels) and the corresponding HOMER motifs outputs (lower panels) in MCF7 and in MDA-MB-231 cells. **(D)** RT-PCR of AS genes in *METTL3* knockdown in the breast cancer cell lines MCF7 and MDA-MB-231. The PSI was calculated in percentage for each gene. Non-specific bands are indicated with an asterisk. *GAPDH* was used as a loading control. **(E)** m^6^A peak distribution in *PHF5A* mRNA in MDA-MB-231 visualized in the Integrative Genomics Viewer (IGV). Input reads are represented in black and the enriched RNA immunoprecipitated in red. The region amplified by qPCR is depicted with a red line below the gene body. **(F)** RT-qPCR of m^6^A RNA immunoprecipitation showing the enrichment of m^6^A in *PHF5A* relative to *GAPDH* in MDA-MB-231. n=3; *** *P-value* < 0.001. **(G)** Venn diagram depicting the overlap of the DSE between *METTL3* knockdown in MDA-MB-231 and *PHF5A* knockout in the breast cancer invasive CA1a cell line (15); *P-value* < 0.05. **(H)** m^6^A peak distribution in *MYC* mRNA in MCF7 (upper panel) and MDA-MB-231 (lower panel) visualized in IGV. Input reads are represented in black and the enriched RNA immunoprecipitated in yellow. The amplified region by qPCR is depicted with a red line below the gene body. **(I)** RT-qPCR of m^6^A RNA immunoprecipitation showing the enrichment of m^6^A in *MYC* relative to *GAPDH* in MCF7 (left panel) and MDA-MB-231 (right panel). n=3; **** *P-value* < 0.0001. **(H)** Venn diagram showing the overlap of the DSE between *METTL3* knockdown in MCF7 and MDA-MB-231 cell lines, and MYC-associated AS events in MDA-MB-436 breast cancer cell line (78); *P-value* < 0.0001.

We then sought to identify whether intronic m^6^A deposition is associated with AS in our breast cancer models. To do so, we performed RT-PCR of nine transcripts harboring intronic m^6^A and two transcripts lacking m^6^A, in MCF7 and MDA-MB-231 upon silencing of *METTL3* (**Figure 3D** and **Supplementary Figure 3F**). We found that METTL3 depletion promoted exon inclusion of the alternative exons of *GNAS, MATR3, POLDIP3* and promoted skipping of the alternative exons of *BAX, COMMD4, MARK3* and the non-m^6^A modified transcripts *FASTK* and *EXOC7* in both breast cancer cell lines. Depletion of METTL3 led to a decrease of the inclusive isoform of *MCM3* in MCF7 whereas no significant difference was observed between the knockdowns and control cells in MDA-MB-231. In contrast, a decrease of the inclusive isoform was observed for *RAP1B* upon knockdown of *METTL3* in MDA-MB-231 in comparison to MCF7, for which there was no change after depletion of METTL3. Altogether, these results show that intronic m^6^A influences splicing.

The expression of splicing factors is generally dysregulated in breast cancer leading to tumor-specific AS events (61-63). Thus, we analyzed the expression of the spliceosome-associated proteins in the SF3A/B and U2AF core complex and key splicing regulators, in control and knockdown of *METTL3* in all our cellular models (**Supplementary Figure 4A**). In the SF3A/B sub-complex, *SF3A3* expression was higher in MDA-MB-231 in agreement with recent observations stating that SF3A3 predicts molecular and phenotypic features of aggressive human breast cancers (22). Likewise, *SF3B4* displayed higher expression levels not only in MDA-MB-231 but also in MCF7 cells. The transcript levels of SR proteins and hnRNPs during breast tumorigenesis were heterogeneous, being some components up-regulated (e.g. *SRSF1, SRSF3, SRSF9*, and *TRA2B*) and other down-regulated (e.g. *SRSF5* and *SRSF8*) in MCF7 and MDA-MB-231 compared to MCF10-A. Additionally, we found that several components of the SF3A/B and U2AF core complex were targets of m^6^A modification, although we did not detect a major effect on METTL3-mediated expression regulation by assessing the RNA steady levels of those spliceosome-associated transcripts (**Supplementary Figure 4A**). This is consistent with GO analysis of our RNA-seq data that did not reveal dysregulation of splicing-associated categories (**Figure 2I-J**).

PHF5A has been recently identified as a critical splicing factor that promotes breast cancer progression (15). We identified *PHF5A* as a potential target of METTL3 harboring m^6^A modification in MDA-MB-231 cell line (**Figure 3E** and **Supplementary Figure 4A**), which was validated by MeRIP followed by RT-qPCR (**Figure 3F**). We further explored the possibility that METTL3 regulates AS indirectly by influencing the expression of PHF5A *via* m^6^A deposition. To this end, we overlapped the DSE dataset from *METTL3* knockdown in MDA-MB-231 with publicly available DSE from a metastatic breast cancer cell line (CA1a) knockout of *PHF5A* (15) (**Figure 3G**). Our analysis revealed that 26.8% of the METTL3-dependent AS switches overlapped with PHF5A-regulated AS, including genes that were validated in our study upon *METTL3* knockdown. Notably, the common DSE were enriched in the biological processes for “cell-cycle”, “cell division”, “spliceosomal snRNP assembly”, “RNA splicing” among others (**Supplementary Figure 4B**). Additionally, terms related to breast cancer such as “mammary neoplasms” and “mammary carcinoma” were enriched in the GO analysis for diseases, highlighting that both METTL3 and PHF5A regulate tumor-associated AS switches (**Supplementary Figure 4B**).

Overexpression or hyperactivation of the transcription factor MYC occurs in most human cancers, and MYC-mediated upregulation of core splicing factors is critical for sustaining growth in MYC-driven tumors (23,64). Previous studies have illustrated that *MYC* mRNA harbors m^6^A modification (65,66). Indeed, this was also the case in both MCF7 and MDA-MB-231 breast cancer cell lines (**Figure 3H-I**). Hence, we sought to study whether METTL3 could regulate AS through MYC. To this end, we overlapped the DSE datasets from knockdown of *METTL3* in MDA-MB-231 and MCF7 with publicly available DSE from MYC-driven AS switches (**Figure 3J**). We found that ∼22% and ∼30% of the METTL3-associated AS events in MCF7 and MDA-MB-231, respectively, overlapped with the MYC-associated AS events (**Supplementary Figure 4C-D**). Similar to the overlap with the PHF5A database, GO analysis of the common AS events in the Disease category revealed an enrichment for the term “breast cancer” (**Supplementary Figure 4C-D**). Additionally, GO for the biological processes category were enriched for terms related to splicing such as “mRNA splicing, *via* spliceosome”, “RNA splicing”, and “mRNA processing”, as well as terms related to “autophagy”, “cell division”, and “regulation of cell cycle”. Overall, these results suggest that METTL3 may indirectly regulate AS in breast cancer *via* m^6^A deposition in splicing factor *PHF5A* and in *MYC* mRNA.

### Identification of breast cancer prognosis-related AS events

To interrogate whether the DSE events that were validated in the MCF7 and MDA-MB-231 cell lines could define a breast cancer-associated AS signature in patients, we analyzed The Cancer Genome Atlas (TCGA) SpliceSeq datasets as well as their associated clinical information for *COMMD4, GNAS, MATR3, RHOC, MARK3, POLDIP3, FASTK, BAX*, and *EXOC7*. For *COMMD4*, two alternative isoforms were analyzed (namely *COMMD4_*AS1 and *COMMD4_*AS2). We observed that the AS events for *COMMD4_*AS2, *GNAS, MATR3* COMMD4_AS1 and *RHOC* displayed a significant higher PSI value in cancer patients than in normal samples (**Figure 4A-E**), while the PSI values were significantly lower for *MARK3, POLDIP3 and FASTK* in patients with breast cancer (**Figure 4F-H**). However, our analysis showed no difference in the PSI values for *BAX*, and *EXOC7* between the cancer patients and normal samples (**Figure 4G-J**). We next employed the same tool to interrogate the PSI values for each DSE mentioned above at different grades of breast cancer (**Supplementary Figure 5A-J**). In comparison to the normal samples (M0), AS switches occurring in patients with breast cancer metastasis (M1) were significantly different in *COMMD4_*AS2, *MARK3*, and *MATR3* (**Supplementary Figure 5A-C**). Inclusions of alternative exons were more prevalent for *COMMD4_*AS2 and *MATR3*, while exclusions were more prevalent for *MARK3. POLDIP3, FASTK, COMMD4*_AS1, *GNAS* and *RHOC* did not show a significant difference between patients with metastasis and normal samples (**Supplementary Figure 5D-H**). However, exclusions of alternative exons were more prevalent for *POLDIP3, FASTK* and *COMMD4*_AS1 in patients with no metastasis (M0) compared to normal samples, whereas inclusions were more prevalent for *GNAS* and *RHOC*. Additionally, no significant difference in the PSI values were found for *BAX* and *EXOC7* in patients compared to the normal samples (**Supplementary Figure 5I-J**).

**Figure 4.**
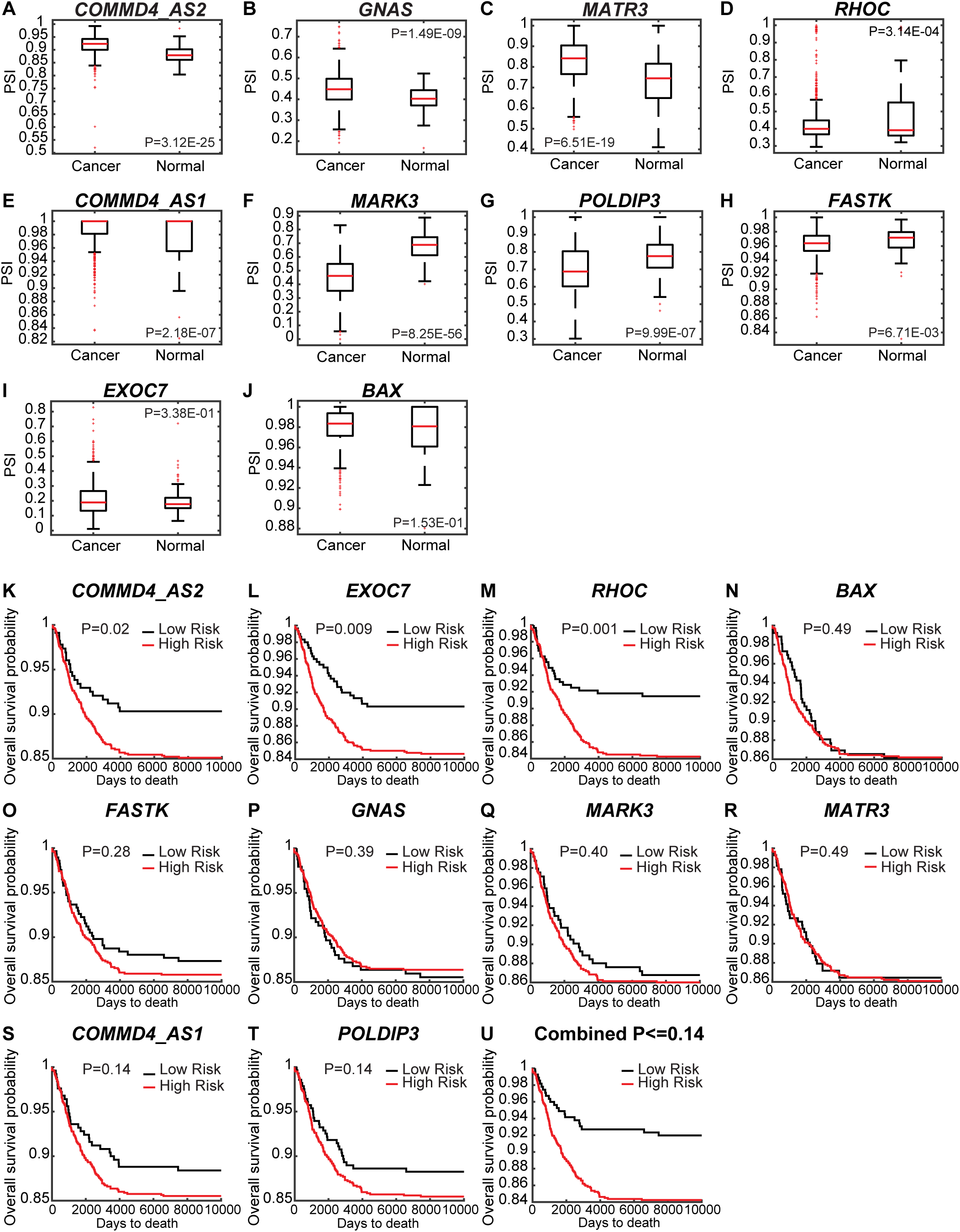
Identification of breast cancer prognosis-related AS events. **(A-J)** PSI values in breast cancer patients (1094 samples) and normal samples (113 samples) for *COMMD4_AS2, GNAS, MATR3, RHOC, COMMD4_AS1, MARK3, POLDIP3, FASTK, EXOC7*, and *BAX*. Data were taken from the TCGA SpliceSeq database. **(K-U)** Kaplan-Meier plots of overall survival for breast cancer patients classified according to the DSE expression (low or high) for *COMMD4_AS2, GNAS, MATR3, RHOC, COMMD4_AS1, MARK3, POLDIP3, FASTK, EXOC7*, and *BAX; P-value* < 0.05.

We next investigated more in depth the AS events within the different stages of breast cancer i.e., from stage I, where the tumor has not spread to lymph nodes or outside the breast, to stage IV in which the cancer has spread to distant organs. Although we found variabilities along the stages, which reflects the heterogeneity of this disease, *COMMD4_*AS2, *MARK3, MATR3, POLDIP3, COMMD4_*AS1, and *GNAS* underwent AS switches in almost all stages of breast cancer, while *RHOC* displayed significant AS switches during stage IIB (**Supplementary Figure 5K-Q**). Nonetheless, *FASTK, BAX* and *EXOC7* did not show a significant difference between patients with breast cancer at different stages and the normal samples (**Supplementary Figure 5R-T**).

We then explored the relationship between DSE and the prognosis of breast cancer patients. To this end, the overall survival rate of breast cancer patients was divided in high or low-risk groups in association to each AS event. Kaplan-Meier curves for *COMMD4_*AS2, *EXOC7* and *RHOC* (*P-value* < 0.05) revealed that patients with a high-risk score had a worse prognosis (**Figure 4K-M**), while the AS events for *BAX, FASTK, GNAS, MARK3*, and *MATR3* were not associated to the survival rate (**Figure 4N-R**). Additionally, *COMMD4_AS1* and *POLDIP3* (**Figure 4S-T**) presented a trend towards a worse prognosis.

Taken together, the AS events analyzed in *COMMD4_AS2, EXOC7, RHOC, COMMD4_*AS1 and *POLDIP3* were related to a worse breast cancer prognosis (**Figure 4U**), and could constitute potential prognosis biomarkers. Furthermore, *COMMD4, MARK3, MATR3, POLDIP3* could be used as biomarkers to specify the stage of the disease.

## DISCUSSION

In the last decade, m^6^A has been established as an important layer of post-transcriptional control of gene expression, and its dysregulated deposition has been defined to be critical for breast cancer initiation, progression and metastasis (43). Although several studies have shown the role of m^6^A in splicing regulation (9,25,38,41,58), to our knowledge, the function of m^6^A in breast cancer-associated AS switches has not been reported yet. Here, we identify genome-wide METTL3-regulated AS events in breast cancer cell lines, and reveal both direct and indirect connections between m^6^A and AS.

We profiled the transcriptome of the normal epithelial MCF10-A, MCF7 and MDA-MB-231 cell lines, with the last two representing distinct breast cancer subtypes. We observed global changes in AS of common transcripts across the three cell lines. Analysis of the AS landscape also revealed a cell-type specific AS signature of a number of genes involved in critical functions for breast tumorigenesis, such as mitotic nuclear division, MAPK signaling cascade, and DNA repair. We validated a selection of AS events, many of them with a known function in migration, invasion and EMT. Interestingly, we found some of these splicing patterns to be similar between the non-tumorigenic MCF10-A and the invasive MDA-MB-231 cell lines. MCF10-A can be grown three-dimensionally (3D) in matrigel mimicking the acinar structure of the mammary gland (67), and opposite splicing patterns between 3D and 2D MCF10-A cultures have been reported (16). Hence, future studies should address which cellular conditions, 3D or 2D, are more faithful to non-tumorigenic epithelial cells. Additionally, a molecular characteristic of MCF10-A includes amplification of *MYC*, which has been reported to play a critical role in oncogenic AS switches (23,64). Nevertheless, the majority of the validated AS events displayed cell-type specific but also common patterns between both breast cancer cell lines.

Although METTL3 has been extensively studied for over a decade, our knowledge about its role in cancer is still limited. Hence, we also interrogated genome-wide METTL3-regulated AS events in breast cancer cell lines. Our data further demonstrates that METTL3 regulates tumor-associated AS switches in breast cancer, and that METTL3 depletion causes mainly alternative first and exon skipping events. It has been shown that the reader YTHDC1 binds to m^6^A sites and recruits the splicing factor SRSF3 to promote exon inclusion (25). Hence, it is plausible that lower m^6^A deposition upon depletion of METTL3 leads to decreased YTHDC1 binding, which in turn promotes exon skipping. Differentially expressed and spliced genes in *METTL3* knockdown MCF7 cells were enriched in cancer-associated categories such as MAPK cascade, cell migration and cell-cell adhesion. However, in MDA-MB-231 we found that most of the biological processes were subjected exclusively to regulation by AS but not by changes in gene expression or *vice versa*. For instance, there was a striking enrichment for splicing-related categories in DSE. Hence, it is likely that the observed METTL3-mediated AS phenotype also results from differential splicing events occurring in transcripts encoding splicing factors, although this hypothesis warrants further study. Nevertheless, our findings highlight that many biological processes occurring in breast cancer cells are regulated only by METTL3-induced AS, expanding the repertoire of functions of METTL3 in tumorigenesis.

Our findings furthermore revealed a significant m^6^A deposition near splice junction sites of mRNAs. This was only true when comparing genome-wide DSE with MeRIP-seq data from the same cell line, emphasizing the notion of a cell-type specific AS and m^6^A signature. Similar to previous observations (41), intronic m^6^A deposition was mostly found in non-DRACH sequences. One limitation of our study is that we used publicly available MeRIP-seq data by which m^6^A regions are detected as the enrichment of immunoprecipitated RNA relative to input RNA. Therefore, the m^6^A site at nucleotide resolution and the m^6^A stoichiometry cannot be interrogated with this conventional antibody-based approach, thereby hampering the identification of direct m^6^A effects on AS. Noteworthy, new technologies such as DART-seq and m^6^A-SAC-seq have identified more m^6^A sites than previously known, yet this data is not available for the cell lines used in our study (68,69). This suggests that the plethora of potential splice sites regulated by m^6^A is underestimated. Additionally, single cell analysis has uncovered substantial heterogeneity of m^6^A sites across individual cells (70). Indeed, many m^6^A sites that are highly methylated at the population level show a low or an absence of methylation in a substantial number of individual cells, which could further impede the assessment of m^6^A-associated DSE. Noteworthy, gene transcripts arising after depletion of METTL3 likely encoded for functional proteins, as we did not observe an enrichment for PTC. Thus, the METTL3-dependent AS switch may generate new isoforms (e.g. *MATR3*) or alter the proportion of existing isoforms (e.g. *BAX*). *EXOC7* AS switches have been previously reported to occur during EMT in breast cancer (71); one isoform containing an alternative 3’
s region of exon 8 (isoform 5 or E) promotes a non-invasive epithelial phenotype, while another isoform lacking this region (isoform 2 or M) has been associated with a mesenchymal aggressive phenotype (71). Herein, we have uncovered a novel *EXOC7* AS switch occurring in breast tumorigenesis. We found that a long isoform *EXOC7-L*, including the exon 7, is only present in MCF7 and MDA-MB-231 cell lines, whereas the short isoform *EXOC7-S*, lacking exon 7, is present in both MCF10-A and MCF7. Such AS switching has been also reported to occur in human fibroblasts (72). Strikingly, our results showed a decrease of exon 7 inclusion of *EXOC7* upon silencing of METTL3. Additionally, we report that this event of EXOC7 isoform switch has a prognosis value for breast cancer patients.

METTL3-regulated AS is not limited to m^6^A deposition at intronic regions. In addition to *cis*-acting RNA elements, dysregulated expression of splicing factors and their mediated splicing events are widely acknowledged to generate distinct AS events. Hence, our analysis revealed differential expression of spliceosome subcomplex components across MCF10-A, MCF7 and MDA-MB-231 cell lines. We further found that several mRNAs encoding splicing factors were decorated with m^6^A modification. Yet, although these transcripts harbor m^6^A, no major change at the mRNA level was found after silencing of *METTL3*. m^6^A modification in mRNA is known to primarily affect export, splicing, RNA stability, and translation, and therefore, m^6^A-mediated control of gene expression might not be reflected by changes in steady-state mRNA levels assessed by RNA-seq. One possible mechanism by which METTL3 can potentially regulate AS is *via* the splicing factor PHF5A and the proto-oncogene MYC as: i) both of them are implicated in AS in breast cancer (15,64); *PHF5A* and *MYC* mRNAs are decorated with the m^6^A mark; and iii) METTL3-mediated DSE significantly overlapped with PHF5A- and MYC-regulated DSE, although we cannot disregard the possibility that those AS events are more sensitive to switch upon perturbation of the splicing factor that regulates them. Nevertheless, further elucidation of METTL3-mediated regulation of PHF5A and MYC expression that led to splicing changes is needed. In addition to these important examples, the potential dysregulation of other splicing factors and their subsequent splicing events still need to be explored, and this information might be particularly valuable to deepen our understanding of the biological relevance of METTL3 in breast cancer.

In this study, we found a higher PSI value in breast cancer patients than in normal samples for *COMMD4_AS2, GNAS, MATR3, RHOC and COMMD4_AS1*, whereas the PSI value was lower for *MARK3, POLDIP3* and FASTK. Nonetheless, we observed no differences in the case of *BAX* and *EXOC7*. These results did not fully reflect our findings from the DSE validated in the non-tumorigenic and the breast cancer cell lines *in vitro*. One possible explanation is that the AS events database gathers information of all breast cancer subtypes and each subtype is associated with a unique AS signature. However, we cannot rule out the possibility that other factors influence the apparent differences between cell lines and patients, a key challenge for translating findings to the clinic. Despite that the AS events analyzed have been previously described in breast cancer and other types of cancer, we have observed variabilities in the change of the PSI value along the progression of breast cancer. For instance, *EXOC7* and *FASTK* displayed a lower PSI at stage III and an increase at stage IV, while for *POLDIP3* the PSI increased at stage III but decreased at stage IV. This indicates that metastasis evolving from a primary tumor is a complex process whereby the tumor acquires metastatic characteristics through additional variables. Additionally, further studies should address whether the difference between our validation *in vitro* and the TCGA SpliceSeq analysis could arise from the cancer heterogeneity in patients, or whether the cancers originate from a single progenitor cell or from polyclonal seeding, leading to different outcomes during tumorigenesis (73,74). Moreover, supporting our results, previous studies have shown genetic differences between primary tumors and lymph node metastases (75-77), because cells can evolve independently of the primary tumor and that different tumor clones can be seeded in parallel to distant sites.

In summary, our study provides further insight into the function of METTL3 and m^6^A in breast cancer by regulating tumor-associated AS switches. Future work should uncover whether these DSE result directly from m^6^A deposition at splice sites or arise from a dysregulated expression of splicing factors, and provide new insights into the regulation and function of m^6^A-associated AS within individual cells from a given population. A better understanding of these molecular mechanisms will then potentially improve the therapeutic opportunities that specifically target breast cancer-associated AS isoforms.

## ACCESSION NUMBERS

All next-generation sequencing data can be publicly accessed in ArrayExpress webserver (E-MTAB-11664).

## SUPPLEMENTARY DATA

Supplementary Data are available at NAR online.

## ACKNOWLEDGEMENT

We would like to thank Aguilo Lab members for useful discussion. We thank Ulf Andersson Vang Ørom for input.

## REFERENCES

1. Kalsotra, A. and Cooper, T.A. (2011) Functional consequences of developmentally regulated alternative splicing. Nat Rev Genet, 12, 715–729.

2. Wang, E.T., Sandberg, R., Luo, S., Khrebtukova, I., Zhang, L., Mayr, C., Kingsmore, S.F., Schroth, G.P. and Burge, C.B. (2008) Alternative isoform regulation in human tissue transcriptomes. Nature, 456, 470–476.

3. Matera, A.G. and Wang, Z. (2014) A day in the life of the spliceosome. Nat Rev Mol Cell Biol, 15, 108–121.

4. Braunschweig, U., Gueroussov, S., Plocik, A.M., Graveley, B.R. and Blencowe, B.J. (2013) Dynamic integration of splicing within gene regulatory pathways. Cell, 152, 1252–1269.

5. Luco, R.F., Allo, M., Schor, I.E., Kornblihtt, A.R. and Misteli, T. (2011) Epigenetics in alternative pre-mRNA splicing. Cell, 144, 16–26.

6. Martin, E., Vivori, C., Rogalska, M., Herrero-Vicente, J. and Valcarcel, J. (2021) Alternative splicing regulation of cell-cycle genes by SPF45/SR140/CHERP complex controls cell proliferation. RNA, 27, 1557–1576.

7. Malla, S., Prasad Bhattarai, D., Groza, P., Melguizo-Sanchis, D., Atanasoai, I., Martinez-Gamero, C., Roman, A.C., Zhu, D., Lee, D.F., Kutter, C. et al. (2022) ZFP207 sustains pluripotency by coordinating OCT4 stability, alternative splicing and RNA export. EMBO Rep, 23, e53191.

8. Paronetto, M.P., Passacantilli, I. and Sette, C. (2016) Alternative splicing and cell survival: from tissue homeostasis to disease. Cell Death Differ, 23, 1919–1929.

9. Jiang, X.L., Liu, B.Y., Nie, Z., Duan, L.C., Xiong, Q.X., Jin, Z.X., Yang, C.P. and Chen, Y.B. (2021) The role of m6A modification in the biological functions and diseases. Signal Transduct Tar, 6.

10. Dvinge, H., Kim, E., Abdel-Wahab, O. and Bradley, R.K. (2016) RNA splicing factors as oncoproteins and tumour suppressors. Nat Rev Cancer, 16, 413–430.

11. Lee, S.C. and Abdel-Wahab, O. (2016) Therapeutic targeting of splicing in cancer. Nat Med, 22, 976–986.

12. Urbanski, L.M., Leclair, N. and Anczukow, O. (2018) Alternative-splicing defects in cancer: Splicing regulators and their downstream targets, guiding the way to novel cancer therapeutics. Wiley Interdiscip Rev RNA, 9, e1476.

13. Park, S., Brugiolo, M., Akerman, M., Das, S., Urbanski, L., Geier, A., Kesarwani, A.K., Fan, M., Leclair, N., Lin, K.T. et al. (2019) Differential Functions of Splicing Factors in Mammary Transformation and Breast Cancer Metastasis. Cell Rep, 29, 2672–2688 e2677.

14. Kahles, A., Lehmann, K.V., Toussaint, N.C., Huser, M., Stark, S.G., Sachsenberg, T., Stegle, O., Kohlbacher, O., Sander, C., Ratsch, G. et al. (2018) Comprehensive Analysis of Alternative Splicing Across Tumors from 8,705 Patients. Cancer Cell, 34, 211-+.

15. Zheng, Y.Z., Xue, M.Z., Shen, H.J., Li, X.G., Ma, D., Gong, Y., Liu, Y.R., Qiao, F., Xie, H.Y., Lian, B. et al. (2018) PHF5A Epigenetically Inhibits Apoptosis to Promote Breast Cancer Progression. Cancer Res, 78, 3190–3206.

16. Anczukow, O., Rosenberg, A.Z., Akerman, M., Das, S., Zhan, L., Karni, R., Muthuswamy, S.K. and Krainer, A.R. (2012) The splicing factor SRSF1 regulates apoptosis and proliferation to promote mammary epithelial cell transformation. Nat Struct Mol Biol, 19, 220–228.

17. Eswaran, J., Horvath, A., Godbole, S., Reddy, S.D., Mudvari, P., Ohshiro, K., Cyanam, D., Nair, S., Fuqua, S.A., Polyak, K. et al. (2013) RNA sequencing of cancer reveals novel splicing alterations. Sci Rep, 3, 1689.

18. Venables, J.P., Klinck, R., Bramard, A., Inkel, L., Dufresne-Martin, G., Koh, C., Gervais-Bird, J., Lapointe, E., Froehlich, U., Durand, M. et al. (2008) Identification of alternative splicing markers for breast cancer. Cancer Res, 68, 9525–9531.

19. Venables, J.P., Klinck, R., Koh, C., Gervais-Bird, J., Bramard, A., Inkel, L., Durand, M., Couture, S., Froehlich, U., Lapointe, E. et al. (2009) Cancer-associated regulation of alternative splicing. Nat Struct Mol Biol, 16, 670–676.

20. Shapiro, I.M., Cheng, A.W., Flytzanis, N.C., Balsamo, M., Condeelis, J.S., Oktay, M.H., Burge, C.B. and Gertler, F.B. (2011) An EMT-driven alternative splicing program occurs in human breast cancer and modulates cellular phenotype. PLoS Genet, 7, e1002218.

21. Anczukow, O., Akerman, M., Clery, A., Wu, J., Shen, C., Shirole, N.H., Raimer, A., Sun, S., Jensen, M.A., Hua, Y. et al. (2015) SRSF1-Regulated Alternative Splicing in Breast Cancer. Mol Cell, 60, 105–117.

22. Ciesla, M., Ngoc, P.C.T., Cordero, E., Martinez, A.S., Morsing, M., Muthukumar, S., Beneventi, G., Madej, M., Munita, R., Jonsson, T. et al. (2021) Oncogenic translation directs spliceosome dynamics revealing an integral role for SF3A3 in breast cancer. Mol Cell, 81, 1453–1468 e1412.

23. Hsu, T.Y., Simon, L.M., Neill, N.J., Marcotte, R., Sayad, A., Bland, C.S., Echeverria, G.V., Sun, T., Kurley, S.J., Tyagi, S. et al. (2015) The spliceosome is a therapeutic vulnerability in MYC-driven cancer. Nature, 525, 384–388.

24. Boccaletto, P., Stefaniak, F., Ray, A., Cappannini, A., Mukherjee, S., Purta, E., Kurkowska, M., Shirvanizadeh, N., Destefanis, E., Groza, P. et al. (2022) MODOMICS: a database of RNA modification pathways. 2021 update. Nucleic Acids Res, 50, D231–D235.

25. Xiao, W., Adhikari, S., Dahal, U., Chen, Y.S., Hao, Y.J., Sun, B.F., Sun, H.Y., Li, A., Ping, X.L., Lai, W.Y. et al. (2016) Nuclear m(6)A Reader YTHDC1 Regulates mRNA Splicing. Mol Cell, 61, 507–519.

26. Roignant, J.Y. and Soller, M. (2017) m(6)A in mRNA: An Ancient Mechanism for Fine-Tuning Gene Expression. Trends Genet, 33, 380–390.

27. Liu, N., Zhou, K.I., Parisien, M., Dai, Q., Diatchenko, L. and Pan, T. (2017) N6-methyladenosine alters RNA structure to regulate binding of a low-complexity protein. Nucleic Acids Res, 45, 6051–6063.

28. Liu, N., Dai, Q., Zheng, G., He, C., Parisien, M. and Pan, T. (2015) N(6)-methyladenosine-dependent RNA structural switches regulate RNA-protein interactions. Nature, 518, 560–564.

29. Liu, J., Yue, Y., Han, D., Wang, X., Fu, Y., Zhang, L., Jia, G., Yu, M., Lu, Z., Deng, X. et al. (2014) A METTL3-METTL14 complex mediates mammalian nuclear RNA N6-adenosine methylation. Nat Chem Biol, 10, 93–95.

30. Ping, X.L., Sun, B.F., Wang, L., Xiao, W., Yang, X., Wang, W.J., Adhikari, S., Shi, Y., Lv, Y., Chen, Y.S. et al. (2014) Mammalian WTAP is a regulatory subunit of the RNA N6-methyladenosine methyltransferase. Cell Res, 24, 177–189.

31. Wang, X., Lu, Z., Gomez, A., Hon, G.C., Yue, Y., Han, D., Fu, Y., Parisien, M., Dai, Q., Jia, G. et al. (2014) N6-methyladenosine-dependent regulation of messenger RNA stability. Nature, 505, 117–120.

32. Haussmann, I.U., Bodi, Z., Sanchez-Moran, E., Mongan, N.P., Archer, N., Fray, R.G. and Soller, M. (2016) m(6)A potentiates Sxl alternative pre-mRNA splicing for robust Drosophila sex determination. Nature, 540, 301–304.

33. Lence, T., Akhtar, J., Bayer, M., Schmid, K., Spindler, L., Ho, C.H., Kreim, N., Andrade-Navarro, M.A., Poeck, B., Helm, M. et al. (2016) m(6)A modulates neuronal functions and sex determination in Drosophila. Nature, 540, 242–247.

34. Kan, L., Grozhik, A.V., Vedanayagam, J., Patil, D.P., Pang, N., Lim, K.S., Huang, Y.C., Joseph, B., Lin, C.J., Despic, V. et al. (2017) The m(6)A pathway facilitates sex determination in Drosophila. Nat Commun, 8, 15737.

35. Liu, X.Y., Li, H.L., Su, J.B., Ding, F.H., Zhao, J.J., Chai, F., Li, Y.X., Cui, S.C., Sun, F.Y., Wu, Z.Y. et al. (2015) Regulation of RAGE splicing by hnRNP A1 and Tra2beta-1 and its potential role in AD pathogenesis. J Neurochem, 133, 187–198.

36. Zhao, Z., Cai, Q., Zhang, P., He, B., Peng, X., Tu, G., Peng, W., Wang, L., Yu, F. and Wang, X. (2021) N6-Methyladenosine RNA Methylation Regulator-Related Alternative Splicing (AS) Gene Signature Predicts Non-Small Cell Lung Cancer Prognosis. Front Mol Biosci, 8, 657087.

37. Huang, X.T., Li, J.H., Zhu, X.X., Huang, C.S., Gao, Z.X., Xu, Q.C., Zhao, W. and Yin, X.Y. (2021) HNRNPC impedes m(6)A-dependent anti-metastatic alternative splicing events in pancreatic ductal adenocarcinoma. Cancer Lett, 518, 196–206.

38. Zhao, X., Yang, Y., Sun, B.F., Shi, Y., Yang, X., Xiao, W., Hao, Y.J., Ping, X.L., Chen, Y.S., Wang, W.J. et al. (2014) FTO-dependent demethylation of N6-methyladenosine regulates mRNA splicing and is required for adipogenesis. Cell Research, 24, 1403–1419.

39. Bartosovic, M., Molares, H.C., Gregorova, P., Hrossova, D., Kudla, G. and Vanacova, S. (2017) N6-methyladenosine demethylase FTO targets pre-mRNAs and regulates alternative splicing and 3’-end processing. Nucleic Acids Res, 45, 11356–11370.

40. Ke, S., Pandya-Jones, A., Saito, Y., Fak, J.J., Vagbo, C.B., Geula, S., Hanna, J.H., Black, D.L., Darnell, J.E., Jr. and Darnell, R.B. (2017) m(6)A mRNA modifications are deposited in nascent pre-mRNA and are not required for splicing but do specify cytoplasmic turnover. Genes Dev, 31, 990–1006.

41. Louloupi, A., Ntini, E., Conrad, T. and Orom, U.A.V. (2018) Transient N-6-Methyladenosine Transcriptome Sequencing Reveals a Regulatory Role of m6A in Splicing Efficiency. Cell Rep, 23, 3429–3437.

42. Zhou, K.I., Shi, H., Lyu, R., Wylder, A.C., Matuszek, Z., Pan, J.N., He, C., Parisien, M. and Pan, T. (2019) Regulation of Co-transcriptional Pre-mRNA Splicing by m(6)A through the Low-Complexity Protein hnRNPG. Mol Cell, 76, 70–81 e79.

43. Kumari, K., Groza, P. and Aguilo, F. (2021) Regulatory roles of RNA modifications in breast cancer. NAR Cancer, 3, zcab036.

44. Destefanis, E., Avsar, G., Groza, P., Romitelli, A., Torrini, S., Pir, P., Conticello, S.G., Aguilo, F. and Dassi, E. (2021) A mark of disease: how mRNA modifications shape genetic and acquired pathologies. RNA, 27, 367–389.

45. Zhao, C., Ling, X., Xia, Y., Yan, B. and Guan, Q. (2021) The m6A methyltransferase METTL3 controls epithelial-mesenchymal transition, migration and invasion of breast cancer through the MALAT1/miR-26b/HMGA2 axis. Cancer Cell Int, 21, 441.

46. Wang, H., Xu, B. and Shi, J. (2020) N6-methyladenosine METTL3 promotes the breast cancer progression via targeting Bcl-2. Gene, 722, 144076.

47. Wu, L., Wu, D., Ning, J., Liu, W. and Zhang, D. (2019) Changes of N6-methyladenosine modulators promote breast cancer progression. BMC Cancer, 19, 326.

48. Bhattarai, D.P. and Aguilo, F. (2022) m(6)A RNA Immunoprecipitation Followed by High-Throughput Sequencing to Map N(6)-Methyladenosine. Methods Mol Biol, 2404, 355–362.

49. Trincado, J.L., Entizne, J.C., Hysenaj, G., Singh, B., Skalic, M., Elliott, D.J. and Eyras, E. (2018) SUPPA2: fast, accurate, and uncertainty-aware differential splicing analysis across multiple conditions. Genome Biology, 19.

50. Anders, S. and Huber, W. (2010) Differential expression analysis for sequence count data. Genome Biol, 11, R106.

51. Benjamini, Y. and Hochberg, Y. (1995) Controlling the False Discovery Rate -a Practical and Powerful Approach to Multiple Testing. J R Stat Soc B, 57, 289–300.

52. Patro, R., Duggal, G., Love, M.I., Irizarry, R.A. and Kingsford, C. (2017) Salmon provides fast and bias-aware quantification of transcript expression. Nat Methods, 14, 417-+.

53. Zhang, Y., Liu, T., Meyer, C.A., Eeckhoute, J., Johnson, D.S., Bernstein, B.E., Nusbaum, C., Myers, R.M., Brown, M., Li, W. et al. (2008) Model-based analysis of ChIP-Seq (MACS). Genome Biol, 9, R137.

54. Quinlan, A.R. and Hall, I.M. (2010) BEDTools: a flexible suite of utilities for comparing genomic features. Bioinformatics, 26, 841–842.

55. Heinz, S., Benner, C., Spann, N., Bertolino, E., Lin, Y.C., Laslo, P., Cheng, J.X., Murre, C., Singh, H. and Glass, C.K. (2010) Simple combinations of lineage-determining transcription factors prime cis-regulatory elements required for macrophage and B cell identities. Mol Cell, 38, 576–589.

56. Ryan, M., Wong, W.C., Brown, R., Akbani, R., Su, X., Broom, B., Melott, J. and Weinstein, J. (2016) TCGASpliceSeq a compendium of alternative mRNA splicing in cancer. Nucleic Acids Res, 44, D1018–1022.

57. Ouyang, D., Yang, P., Cai, J., Sun, S. and Wang, Z. (2020) Comprehensive analysis of prognostic alternative splicing signature in cervical cancer. Cancer Cell Int, 20, 221.

58. Adhikari, S., Xiao, W., Zhao, Y.L. and Yang, Y.G. (2016) m(6)A: Signaling for mRNA splicing. Rna Biol, 13, 756–759.

59. Lee, J.H., Wang, R.Y., Xiong, F., Krakowiak, J., Liao, Z., Nguyen, P.T., Moroz-Omori, E.V., Shao, J.F., Zhu, X.Y., Bolt, M.J. et al. (2021) Enhancer RNA m6A methylation facilitates transcriptional condensate formation and gene activation. Molecular Cell, 81, 3368-+.

60. Li, F., Yi, Y., Miao, Y., Long, W., Long, T., Chen, S., Cheng, W., Zou, C., Zheng, Y., Wu, X. et al. (2019) N(6)-Methyladenosine Modulates Nonsense-Mediated mRNA Decay in Human Glioblastoma. Cancer Res, 79, 5785–5798.

61. Sveen, A., Kilpinen, S., Ruusulehto, A., Lothe, R.A. and Skotheim, R.I. (2016) Aberrant RNA splicing in cancer; expression changes and driver mutations of splicing factor genes. Oncogene, 35, 2413–2427.

62. Silipo, M., Gautrey, H. and Tyson-Capper, A. (2015) Deregulation of splicing factors and breast cancer development. J Mol Cell Biol, 7, 388–401.

63. Wang, Y., Chen, D., Qian, H.L., Tsai, Y.H.S., Shao, S.J., Liu, Q.T., Dominguez, D. and Wang, Z.F. (2014) The Splicing Factor RBM4 Controls Apoptosis, Proliferation, and Migration to Suppress Tumor Progression. Cancer Cell, 26, 374–389.

64. Anczukow, O. and Krainer, A.R. (2015) The spliceosome, a potential Achilles heel of MYC-driven tumors. Genome Med, 7, 107.

65. Aguilo, F., Zhang, F., Sancho, A., Fidalgo, M., Di Cecilia, S., Vashisht, A., Lee, D.F., Chen, C.H., Rengasamy, M., Andino, B. et al. (2015) Coordination of m(6)A mRNA Methylation and Gene Transcription by ZFP217 Regulates Pluripotency and Reprogramming. Cell Stem Cell, 17, 689–704.

66. Batista, P.J., Molinie, B., Wang, J.K., Qu, K., Zhang, J.J., Li, L.J., Bouley, D.M., Lujan, E., Haddad, B., Daneshvar, K. et al. (2014) m(6)A RNA Modification Controls Cell Fate Transition in Mammalian Embryonic Stem Cells. Cell Stem Cell, 15, 707–719.

67. Soule, H.D., Maloney, T.M., Wolman, S.R., Peterson, W.D., Brenz, R., Mcgrath, C.M., Russo, J., Pauley, R.J., Jones, R.F. and Brooks, S.C. (1990) Isolation and Characterization of a Spontaneously Immortalized Human Breast Epithelial-Cell Line, Mcf-10. Cancer Research, 50, 6075–6086.

68. Hu, L., Liu, S., Peng, Y., Ge, R., Su, R., Senevirathne, C., Harada, B.T., Dai, Q., Wei, J., Zhang, L. et al. (2022) m(6)A RNA modifications are measured at single-base resolution across the mammalian transcriptome. Nat Biotechnol.

69. Meyer, K.D. (2019) DART-seq: an antibody-free method for global m(6)A detection. Nat Methods, 16, 1275–1280.

70. Tegowski, M., Flamand, M.N. and Meyer, K.D. (2022) scDART-seq reveals distinct m(6)A signatures and mRNA methylation heterogeneity in single cells. Mol Cell, 82, 868–878 e810.

71. Lu, H.Z., Liu, J.L., Liu, S.J., Zeng, J.W., Ding, D.Q., Carstens, R.P., Cong, Y.S., Xu, X.W. and Guo, W. (2013) Exo70 Isoform Switching upon Epithelial-Mesenchymal Transition Mediates Cancer Cell Invasion. Dev Cell, 27, 560–573.

72. Georgilis, A., Klotz, S., Hanley, C.J., Herranz, N., Weirich, B., Morancho, B., Leote, A.C., D’Artista, L., Gallage, S., Seehawer, M. et al. (2018) PTBP1-Mediated Alternative Splicing Regulates the Inflammatory Secretome and the Pro-tumorigenic Effects of Senescent Cells. Cancer Cell, 34, 85-+.

73. Navin, N., Kendall, J., Troge, J., Andrews, P., Rodgers, L., McIndoo, J., Cook, K., Stepansky, A., Levy, D., Esposito, D. et al. (2011) Tumour evolution inferred by single-cell sequencing. Nature, 472, 90–U119.

74. Nowell, P.C. (1976) Clonal Evolution of Tumor-Cell Populations. Science, 194, 23–28.

75. Schmidt-Kittler, O., Ragg, T., Daskalakis, A., Granzow, M., Ahr, A., Blankenstein, T.J.F., Kaufmann, M., Diebold, J., Arnholdt, H., Muller, P. et al. (2003) From latent disseminated cells to overt metastasis: Genetic analysis of systemic breast cancer progression. P Natl Acad Sci USA, 100, 7737–7742.

76. Torres, L., Ribeiro, F.R., Pandis, N., Andersen, J.A., Heim, S. and Teixeira, M.R. (2007) Intratumor genomic heterogeneity in breast cancer with clonal divergence between primary carcinomas and lymph node metastases. Breast Cancer Res Tr, 102, 143–155.

77. Vecchi, M., Confalonieri, S., Nuciforo, P., Vigano, M.A., Capra, M., Bianchi, M., Nicosia, D., Bianchi, F., Galimberti, V., Viale, G. et al. (2008) Breast cancer metastases are molecularly distinct from their primary tumors. Oncogene, 27, 2148–2158.

78. Wu, S.Y., Xiao, Y., Wei, J.L., Xu, X.E., Jin, X., Hu, X., Li, D.Q., Jiang, Y.Z. and Shao, Z.M. (2021) MYC suppresses STING-dependent innate immunity by transcriptionally upregulating DNMT1 in triple-negative breast cancer. J Immunother Cancer, 9.

