## Supplementary Information for "METTL3 regulates breast cancer-associated alternative splicing switches"

#### Index of Supplemental Figures:


- **Supplemental Figure 1.** Identification of DSE in breast cancer cell lines.
- **Supplemental Figure 2.** Genome-wide analysis of METTL3-mediated AS.
- **Supplemental Figure 3.** METTL3 mediates AS *via* $m^6A$ deposition.
- **Supplemental Figure 4.** $m^6A$ regulates PHF5A- and MYC-associated AS events.
- **Supplemental Figure 5.** DSE signature in breast cancer patients.


#### Index of Supplemental Tables:


- **Supplemental Table 1.** Primers and shRNAs sequences used in this study.
- **Supplemental Table 3.** Function in cancer and type of altered AS event for the transcripts assessed in this study.


#### Index of Supplemental Tables (provided as a single Excel spreadsheet):


- **Supplemental Table 2.** List of the DSE in the breast cancer cell lines MCF7 and MDA-MB-231 compared to the non-tumorigenic cell line MCF10-A.
- **Supplemental Table 4.** List of the DSE upon *METTL3* knockdown in MCF7 and MDA-MB-231 cell lines.

**Supplemental Figure 1**

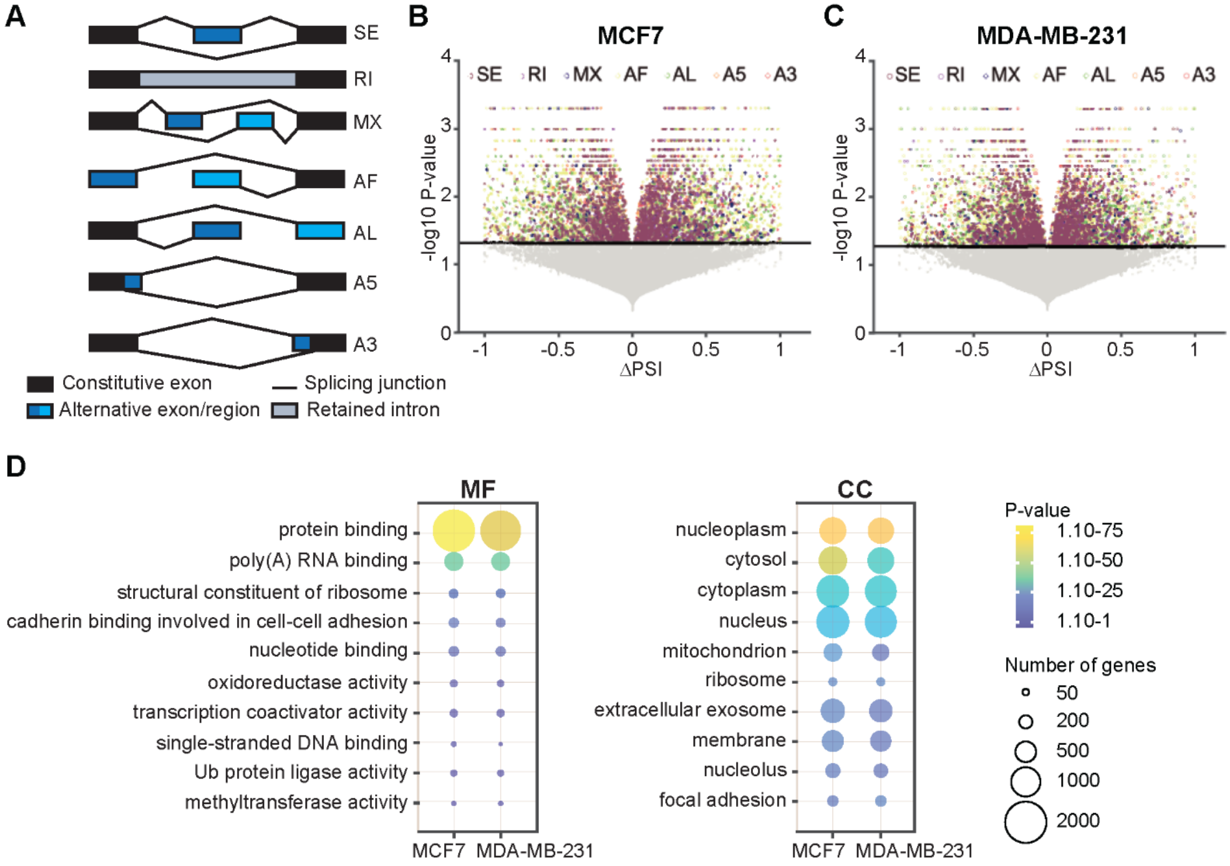


**Supplemental Figure 1.** Identification of differential AS events (DSE) in breast cancer cell lines. **(A)** Schematics of the seven types of AS events. SE: skipped exon; RI: retained intron; MX: mutually exclusive exon; AF: alternative first exon; AL: alternative last exon; A5: 5´ splice site; A3: 3´ splice site. **(B-C)** Volcano plots representing the ΔPSI of the differentially spliced genes in MCF7 and MDA-MB-231 related to MCF10-A cells. The significant DSE are shown in a color code. The y-axis represents the –log10 *P-value* (*P-value* < 0.05). **(C)** Dot plots representing the Gene Ontology (GO) enrichment analysis of the common AS genes in MCF7 and MDA-MB-231. CC: cellular component, MF: molecular function. The size and the color of the dots are proportional to the number of genes enriched in each GO term and the significance of the enrichment ($1.10^{-75}$ < *P-value* < $1.10^{-1}$), respectively.

**Supplemental Figure 2**

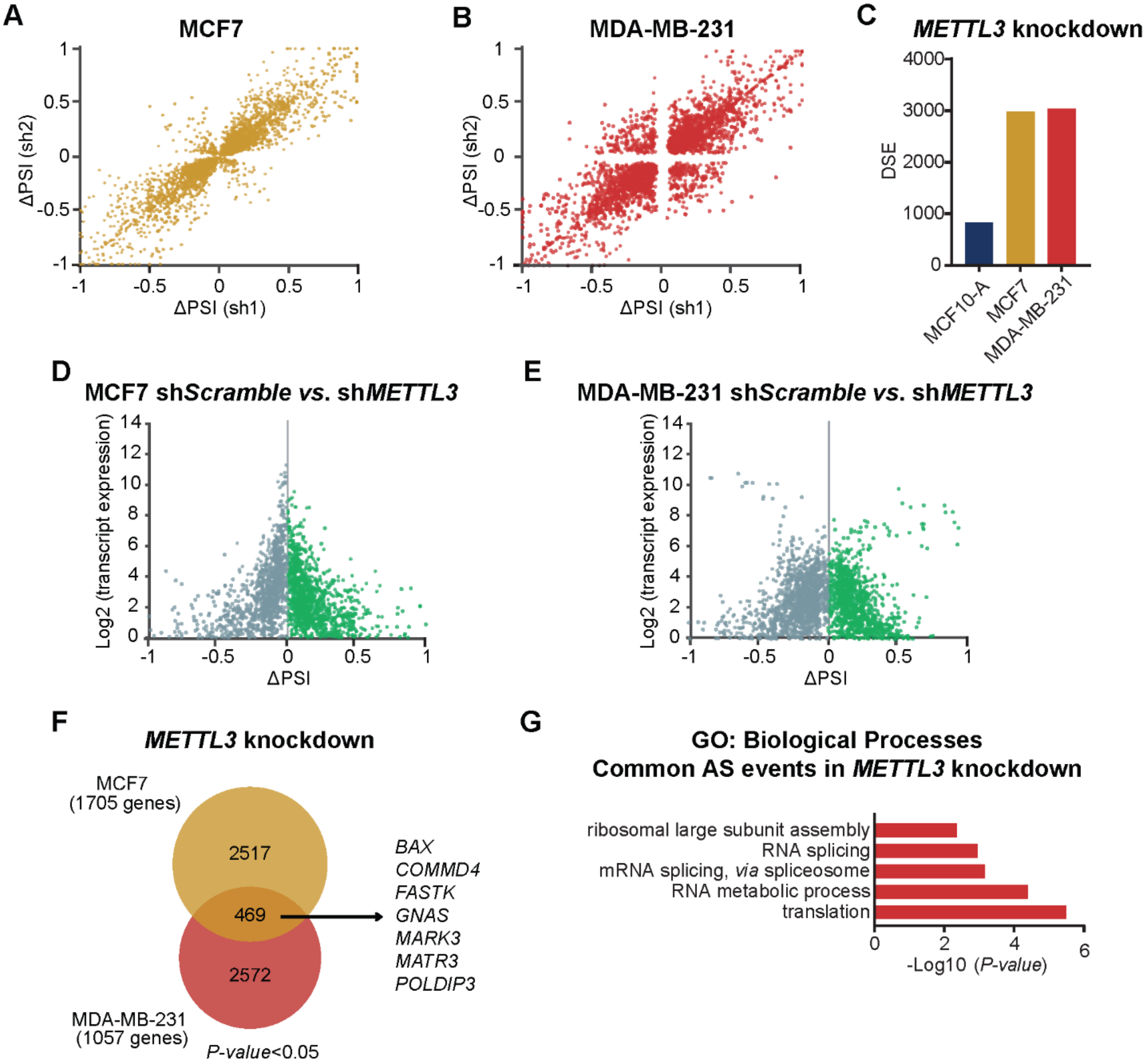


**Supplemental Figure 2.** Genome-wide analysis of METTL3-mediated AS. **(A-B)** Correlation of ΔPSI between biological replicates upon silencing of *METTL3* (sh1 and sh2) in MCF7 and in MDA-MB-231 cells. **(C)** Number of DSE in the non-tumorigenic cell line MCF10-A and the breast cancer cell lines MCF7 and MDA-MB-231 upon silencing of *METTL3*. **(D-E)** Volcano plot showing the correlation between gene expression levels and ΔPSI resulted from RNA-seq data analysis in control and METTL3 depleted MCF7 and MDA-MB-231 cell lines. **(F)** Venn diagrams depicting the common DSE between MCF7 (yellow) and MDA-MB-231 (red) upon depletion of METTL3. The number of DSE for each cell line is indicated in brackets; *P-value* < 0.05. **(G)** GO analysis of common AS genes between MCF7 and MDA-MB-231 upon depletion of METTL3.

### Supplemental Figure 3

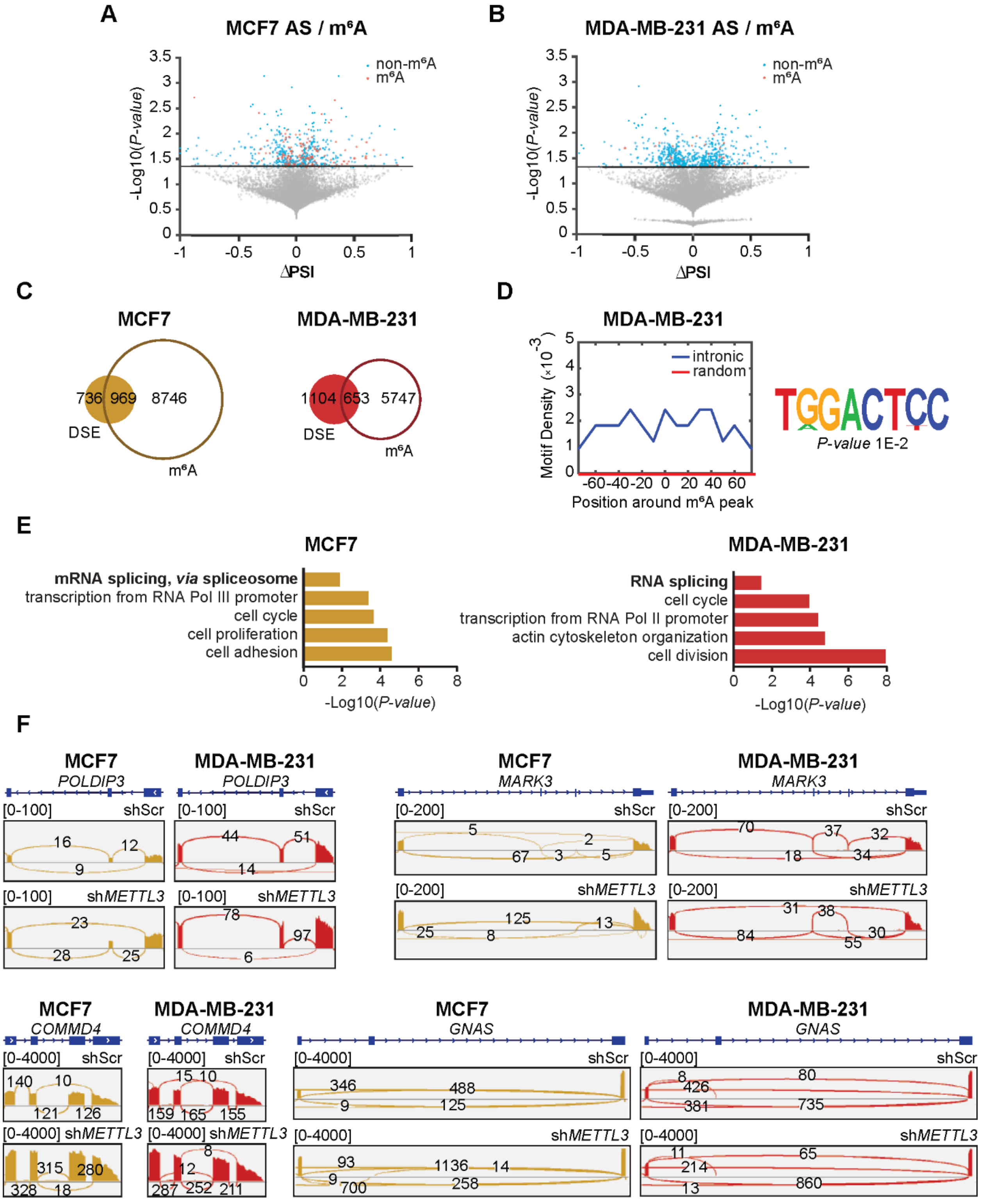


*Figure legend in the next page*

**Supplemental Figure 3.** METTL3 mediates AS *via* $m^6A$ deposition. **(A-B)** Volcano plot of the ∆PSI of the differentially spliced transcripts in MCF7 and in MDA-MB-231. Highlighted in red are the transcripts harboring $m^6A$ and in blue the non-$m^6A$ modified transcripts (*P-value* < 0.05). Grey dots indicate non-significant DSE. Datasets from (1); datasets GEO accession number for MCF7: GSE143441 and MDA-MB-231: GSM5616175. **(C)** Venn diagrams depicting the DSE harboring $m^6A$ mark in knockdown of *METTL3* in MCF7 and MDA-MB-231 cell line. **(D)** DRACH motif density (lowly represented; *P-value* 0.01) of $m^6A$ peaks in the –80 to +80 nt region around the $m^6A$ peak in intronic or random regions (left panel) and the corresponding HOMER motifs outputs (right panel) in MDA-MB-231. **(E)** GO analysis of biological processes associated to the $m^6A$-modified transcripts in the breast tumorigenic cell lines MCF7 and MDA-MB-231 upon depletion of METTL3. *P-value* < 0.05. **(F)** Examples of sashimi plots showing changes of AS events upon knockdown of *METTL3* in MCF7 and MDA-MB-231.

### Supplemental Figure 4

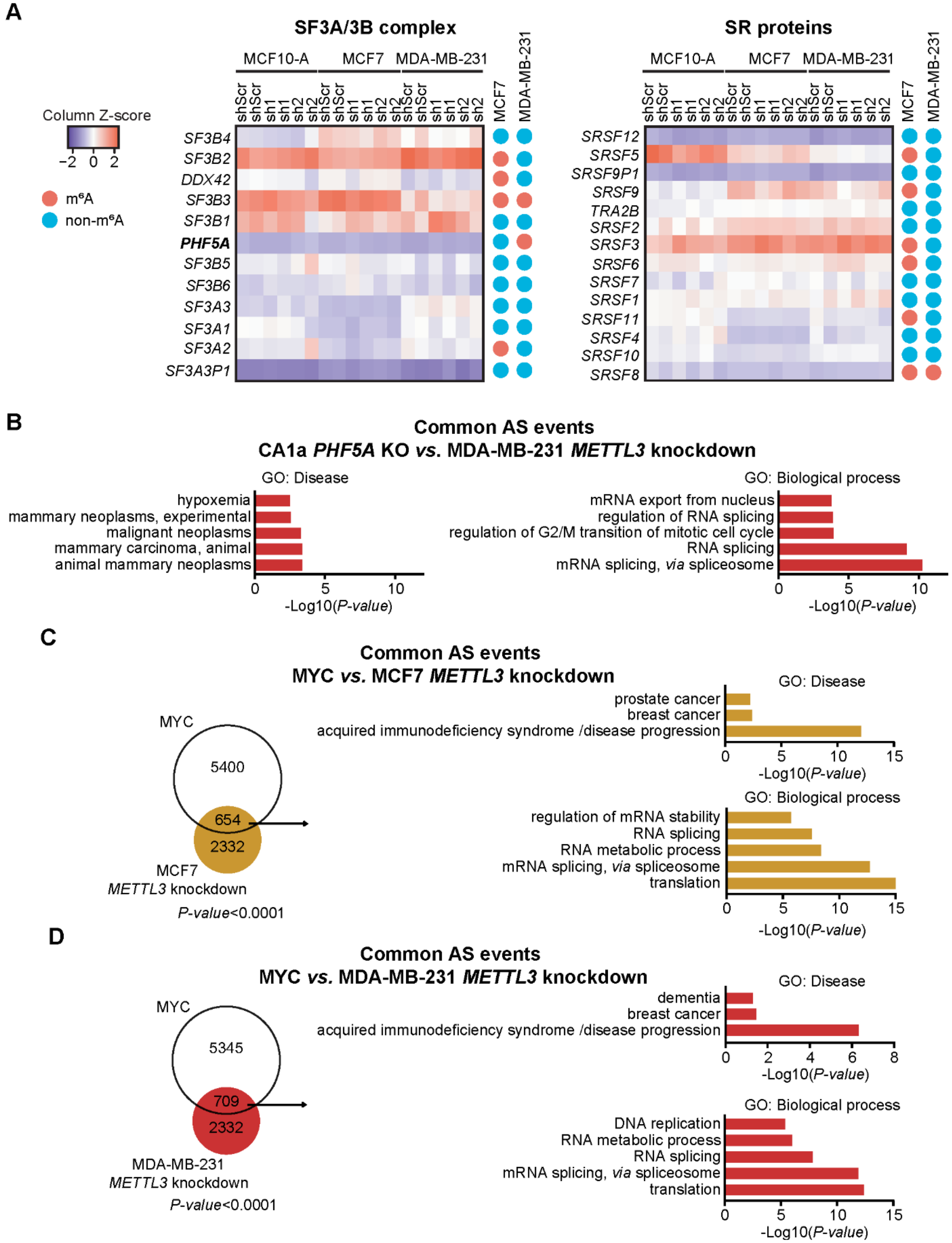


*Figure legend in the next page*

**Supplemental Figure 4.** $m^6A$ regulates PHF5A- and MYC-associated AS events. **(A)** Heatmaps showing the differential expression for splicing factors within the SF3A/3B complex (left panel) and SR proteins (right panel) in MCF10-A, MCF7 and MDA-MB-231 cell lines. For each transcript encoding for a splicing factor is indicated whether it is $m^6A$ modified (red dot) or non-$m^6A$ modified (blue dot). Both heatmaps were scaled by row. The heatmaps were scaled with Z-Score using the $Log_2$(FPKM) expression. **(B)** GO analysis of common AS genes between knockdown of *METTL3* in MDA-MB-231 and knockout of *PHF5A* in CA1a cell line. **(C)** Overlaps between AS events of knockdown of *METTL3* in MCF7 and MYC-associated AS events (left panel); *P-value* < 0.0001. GO analysis of the common genes between AS events between knockdown of *METTL3* in MCF7 and MYC-associated AS events (right panel); *P-value* < 0.05. **(D)** Overlaps between AS events of knockdown of *METTL3* in MDA-MB-231 and MYC-associated AS events (left panel); *P-value* < 0.0001. GO analysis of the common genes between AS events in knockdown of *METTL3* in MDA-MB-231 and MYC-associated AS events (right panel); *P-value* < 0.05.

### Supplemental Figure 5

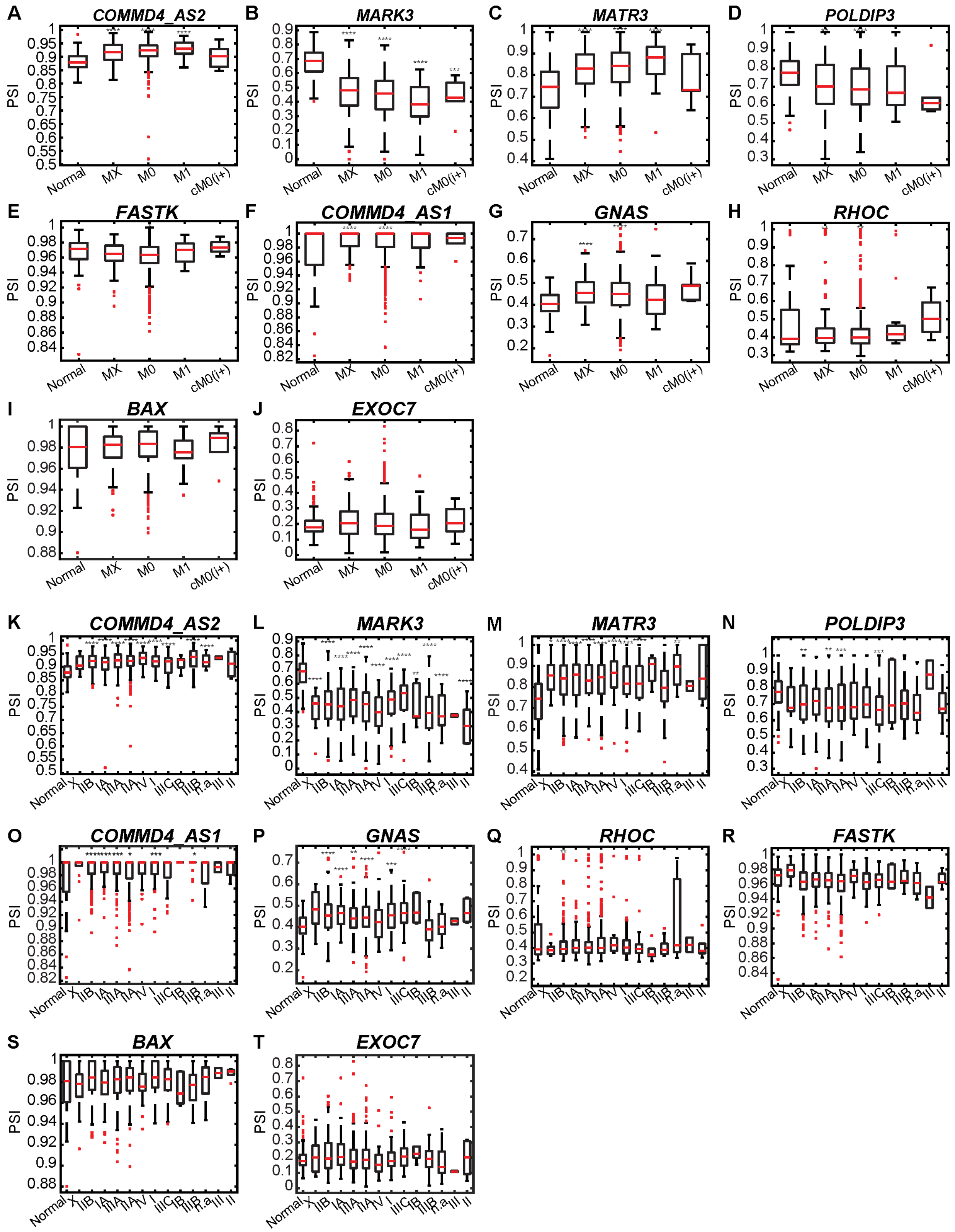


*Figure legend in the next page*

**Supplemental Figure 5.** DSE signature in breast cancer patients. **(A-J)** Box plots representative of the PSI values across breast tumor grades and in normal samples. M0: no distant metastasis, M1: distant metastasis, MX: distant metastasis cannot be assessed, cM0(i+): cancer cells present in blood or bone marrow or in lymph nodes farther from the primary site. **** *P-value* < 0.0001, *** *P-value* < 0.001, ** *P-value* < 0.01; * *P-value* < 0.05. **(K-T)** Box plots representative of the PSI values across breast tumor stages and in normal samples. Stage I: 90 patients; stage IA: 86; stage IB: 6; stage II: 6; stage IIA: 358; stage IIB: 255; stage III: 2; stage IIIA: 154; stage IIIB: 27; stage IIIC: 65; stage IV: 20; stage X: 13; n.a. (no stage assessed): 11; **** *P-value* < 0.0001, *** *P-value* < 0.001, ** *P-value* < 0.01; * *P-value* < 0.05.

**Supplemental Table 1.** Primers and shRNAs sequences used in this study.

| **RT-PCR** | | |
|---|---|---|
| **Gene** | **Forward primer 5´→ 3´** | **Reverse primer 5´→ 3´** |
| *ARHGEF11* | GGCAGCAGGAGGTTACAAAG | TGAGCCTGTTGAGCTTGAGA |
| *BAX* | TGATGGACGGGTCCG | GGCAAAGTAGAAAAGGGCGAC |
| *COMMD4* | GCGAATCCTTGTCCAGTGAA | TCCCGAAGAGCAGTTCAGAG |
| *EBPL* | TTTCCTCTGCCGCATGGT | AACAATGCCAGAGACCCATC |
| *EXOC7* | TGGCCGCAACCAAGATTTCATG | TCGGACAGGTGCTTAACTCGGAAAT |
| *FASTK* | CATCTTGATGTCACTGTGCCA | CAGCAGGGAGAGGTAGCG |
| *GNAS* | AAAAGCACCATTGTGAAGCA | TCAATCGCCTCTTTCAGGTT |
| *KMT5B* | CGGTTTTCGTACGGGCGG | AAATAGTGCCGTGCCCATTC |
| *MARK3* | AAGAGGCACTGCCAGTCGTA | GAGCCCTCATATCTCCCGTTC |
| *MATR3* | GCCTTCACCTGAATGACATCT | GACGACTGTGACTTGCTCCA |
| *MCM3* | TCCAAAGACGGCAGACTCAC | TCCTGCATCTTGCTCAGAGC |
| *POLDIP3* | TGCCTTCATAAACCCACCCA | CATGTGGTGGAGAAAGCCG |
| *RAP1B* | ACATCGCCAAACCTCGCC | CGTTCCTGCAGTATCCAAGA |
| *RHOC* | TCTGAGCCTCCGGCACC | GGACGTAGACCTCCGGAAAC |

| **qPCR** | | |
|---|---|---|
| **Gene** | **Forward primer 5´→ 3´** | **Reverse primer 5´→ 3´** |
| *B-ACTIN* | AGATCAAGGTGGGTGTCTTTC | AGCAATGATCTGAGGAGGGAAG |
| *GAPDH* | TGGTATCGTGGAAGGACTCA | CCAGTAGAGGCAGGGATGAT |
| *METTL3* | AACTGCAACGCATCATTCGG | TTGACACCAACCAAGCAGTG |
| *MYC* | CATCAGCACAACTACGCAGC | GCTGGTGCATTTTCGGTTGT |
| *PHF5A* | TGCAGCTGCCAGAAAACATG | TTTTCCATCCCTACCACGTGTC |

| **shRNAs** | | |
|---|---|---|
| **Target Gene** | **Plasmid** | **Sequence 5´→ 3´** |
| Scramble | pLKO.1-puro-shScramble | CAACAAGATGAAGAGCACCAA |
| *METTL3* | pLKO.1-puro-sh*METTL3*_1 | GCAAGTATGTTCACTATGAAA |
| | pLKO.1-puro-sh*METTL3*_2 | CGTCAGTATCTTGGGCAAGTT |

**Supplemental Table 3.** Function in cancer and type of altered AS event for the transcripts assessed in this study.

| Gene name | Ensembl ID | Function | Type of AS events | Publication |
|---|---|---|---|---|
| *ARHGEF11* | **ENSG00000132694** | EMT | SE exon 38 | (2) |
| | | | SE | (3) |
| | | | A3'SS, RI, A5'SS, SE | (4) |
| | | | SE | (5) |
| | | | SE | (6) |
| *BAX* | **ENSG00000087088** | Apoptosis | | (7) |
| | | | A3'SS,RI | (4) |
| | | | SE | (8) |
| | | | SE | (3) |
| *COMMD4* | **ENSG00000140365** | Unknown | RI, SE | (4) |
| *EBPL* | **ENSG00000123179** | Unknown | MXE | (6) |
| *EXOC7* | **ENSG00000182473** | EMT, invasion | SE | (9) |
| | | | SE | (3) |
| | | | RI,A3'SS, A5'SS, SE | (4) |
| | | | MXE | (6) |
| *FASTK* | **ENSG00000164896** | Apoptosis | RI, A3'SS | (4) |
| | | | RI intron 5 | (10) |
| *GNAS* | **ENSG00000087460** | Signaling, proliferation, migration (EMT) | SE | (8) |
| | | | SE | (4) |
| *KMT5B* | **ENSG00000110066** | Migration, adhesion | SE exon 3, A5'SS, RI | (4) |
| *MARK3* | **ENSG00000075413** | Unknown | SE, RI | (4) |
| | | | SE exon 27 | (8) |
| | | | SE | (6) |
| *MATR3* | **ENSG00000015479** | EMT, apoptosis | | (10) |
| | | | SE, A3'SS, A5'SS | (4) |
| | | | SE | (8) |
| | | | SE | (6) |
| *MCM3* | **ENSG00000112118** | Proliferation, drug resistance | SE | (4) |
| *POLDIP3* | **ENSG00000100227** | Unknown | SE exon 3 | (10) |
| *RAP1B* | **ENSG00000127314** | Cell-cell adhesion | A3'SS, MXE, SE, A5'SS | (4) |
| *RHOC* | **ENSG00000155366** | Cell-cell junction, interferon signaling | SE | (8) |
| | | | A5'SS, A3'SS | (4) |

### LITERATURE CITED


1. Lee, J.H., Wang, R., Xiong, F., Krakowiak, J., Liao, Z., Nguyen, P.T., Moroz-Omori, E.V., Shao, J., Zhu, X., Bolt, M.J. *et al.* (2021) Enhancer RNA m6A methylation facilitates transcriptional condensate formation and gene activation. *Mol Cell*, **81**, 3368-3385 e3369.
2. Itoh, M., Radisky, D.C., Hashiguchi, M. and Sugimoto, H. (2017) The exon 38-containing ARHGEF11 splice isoform is differentially expressed and is required for migration and growth in invasive breast cancer cells. *Oncotarget*, **8**, 92157-92170.
3. Anczukow, O., Akerman, M., Clery, A., Wu, J., Shen, C., Shirole, N.H., Raimer, A., Sun, S., Jensen, M.A., Hua, Y. *et al.* (2015) SRSF1-Regulated Alternative Splicing in Breast Cancer. *Mol Cell*, **60**, 105-117.
4. Park, S., Brugiolo, M., Akerman, M., Das, S., Urbanski, L., Geier, A., Kesarwani, A.K., Fan, M., Leclair, N., Lin, K.T. *et al.* (2019) Differential Functions of Splicing Factors in Mammary Transformation and Breast Cancer Metastasis. *Cell Rep*, **29**, 2672-2688 e2677.
5. Gokmen-Polar, Y., Gu, Y., Gu, X. and Badve, S.S. (2019) Splicing factor ESRP1 controls ER-positive breast cancer progression by altering metabolic pathway genes. *Cancer Research*, **79**.
6. Shapiro, I.M., Cheng, A.W., Flytzanis, N.C., Balsamo, M., Condeelis, J.S., Oktay, M.H., Burge, C.B. and Gertler, F.B. (2011) An EMT-driven alternative splicing program occurs in human breast cancer and modulates cellular phenotype. *PLoS Genet*, **7**, e1002218.
7. Kholoussi, N.M., El-Nabi, S.E.H., Esmaiel, N.N., Abd El-Bary, N.M. and El-Kased, A.F. (2014) Evaluation of Bax and Bak Gene Mutations and Expression in Breast Cancer. *Biomed Res Int*, **2014**.
8. Oh, J., Pradella, D., Shao, C.W., Li, H.R., Choi, N., Ha, J., Ruggiero, S., Fu, X.D., Zheng, X., Ghigna, C. *et al.* (2021) Widespread Alternative Splicing Changes in Metastatic Breast Cancer Cells. *Cells-Basel*, **10**.
9. Lu, H.Z., Liu, J.L., Liu, S.J., Zeng, J.W., Ding, D.Q., Carstens, R.P., Cong, Y.S., Xu, X.W. and Guo, W. (2013) Exo70 Isoform Switching upon Epithelial-Mesenchymal Transition Mediates Cancer Cell Invasion. *Dev Cell*, **27**, 560-573.
10. Zheng, Y.Z., Xue, M.Z., Shen, H.J., Li, X.G., Ma, D., Gong, Y., Liu, Y.R., Qiao, F., Xie, H.Y., Lian, B. *et al.* (2018) PHF5A Epigenetically Inhibits Apoptosis to Promote Breast Cancer Progression. *Cancer Res*, **78**, 3190-3206.
